# Tumor sialylation controls effective anti-cancer immunity in breast cancer

**DOI:** 10.1101/2023.09.20.558571

**Authors:** Stefan Mereiter, Gustav Jonsson, Tiago Oliveira, Johannes Helm, David Hoffmann, Markus Abeln, Ann-Kristin Jochum, Wolfram Jochum, Max J. Kellner, Marek Feith, Vanessa Tkalec, Karolina Wasilewska, Jie Jiao, Lukas Emsenhuber, Felix Holstein, Anna C. Obenauf, Leonardo Lordello, Jean-Yves Scoazec, Guido Kroemer, Laurence Zitvogel, Omar Hasan Ali, Lukas Flatz, Rita Gerardy-Schahn, Anja Münster-Kühnel, Johannes Stadlmann, Josef M. Penninger

## Abstract

Breast cancer is the most common cancer among women. However, the use of immune checkpoint inhibitors, that have revolutionized treatment of multiple cancers, unfortunately remain largely ineffective in most breast cancer patients. Here, we report the most comprehensive glycoproteome map in breast tumor cells, pointing to a key role of sialic acid modifications in mammary cancer. Genetic and pharmacologic inhibition of sialylation repolarizes the tumor microenvironment, leading to a reduction in myeloid-derived suppressor cells and a significant increase in Tcf7^+^ memory and CD8^+^ effector T cells. Mechanistically, sialylation controls cell surface expression of MHC class I and PD-1-ligand on the tumor cells. Functionally, in vivo interference with sialylation on breast cancer cells licenses CD8^+^ T cells to effectively kill the tumors. In multiple immunotherapy-resistant breast tumor models, we also show that the abrogation of sialylation sensitizes to anti-PD-1 immune checkpoint therapy. We further demonstrate that hyper-sialylation occurs in over half of human breast cancers tested and correlates with poor T cell infiltration. Our results establish sialylation as a central immunoregulator in breast cancer, orchestrating multiple pathways of immune evasion. Targeting tumor sialylation licenses immunologically inert mammary tumors to be efficiently eliminated by anti-cancer immunity and sensitizes to immune checkpoint therapy.

## Introduction

Immune checkpoint inhibitors (ICIs) have rapidly become a standard of care for multiple cancers, harnessing the patient’s immune system to combat the disease. However, the currently approved ICIs have shown limited efficacy in breast cancer (BC) (*1, 2*). Nevertheless, BCs often exhibit abundant tumor-infiltrating lymphocytes, albeit rarely expressing programmed death-ligand 1 (PD-L1) (*3*). This suggests that alternative pathways of immunoregulation are in place that could be exploited for BC immunotherapy.

Sialylation, the addition of sialic acid to glycoproteins, glycolipids, or glyco-RNA, is altered in multiple cancers (*4*). Sialic acids, negatively charged nine-carbon sugars that cap glycan structures, play an important role in various cell-cell and cell-matrix interactions (*5*). In recent years, sialylation has garnered attention due to its emerging importance in immunoregulation including anti-tumor immune responses (*6, 7*). Sialylation is therefore a promising candidate as a targetable immunoregulatory pathway. Despite this promise, our current understanding of the role of sialylation in breast cancer and whether sialylation might be mechanistically involved in the resistance of breast cancer to ICI is largely incomplete.

Employing genetic and pharmacological analyses in murine models, human breast cancer cell lines, and clinical specimens, we show that alterations in sialylation enhance tumor recognition and license mammary cancer treatment to effective PD-1 inhibition, turning immunological dormant breast tumors into immune responsive and druggable tumors.

## Results

### Functional and structural glycan characterization of the murine breast cancer models

The surface glycome of cells is the result of the complex interplay of numerous glycosyltransferases. Mouse and human genomes harbor each over 200 of these transferase genes, and their activity is influenced by factors such as the condition of the secretory pathway, the metabolic state of the cells, and the overall proteome composition (*8*). As a result, the glycome of cancer cells cannot be reliably inferred from gene expression data sets. We therefore conducted comprehensive glycome analyses on three commonly used murine breast cancer cell line models 4T1, EMT6, and E0771. Using cell surface glycan flow cytometric analysis employing a panel of plant lectins with well-defined specificities and structural N-glycomic analysis using mass spectrometry (Fig. 1A and Suppl Fig. 1A), we consistently observed that the 4T1 cells exhibited the highest levels of sialylation among the cell lines tested.

**Figure 1.**
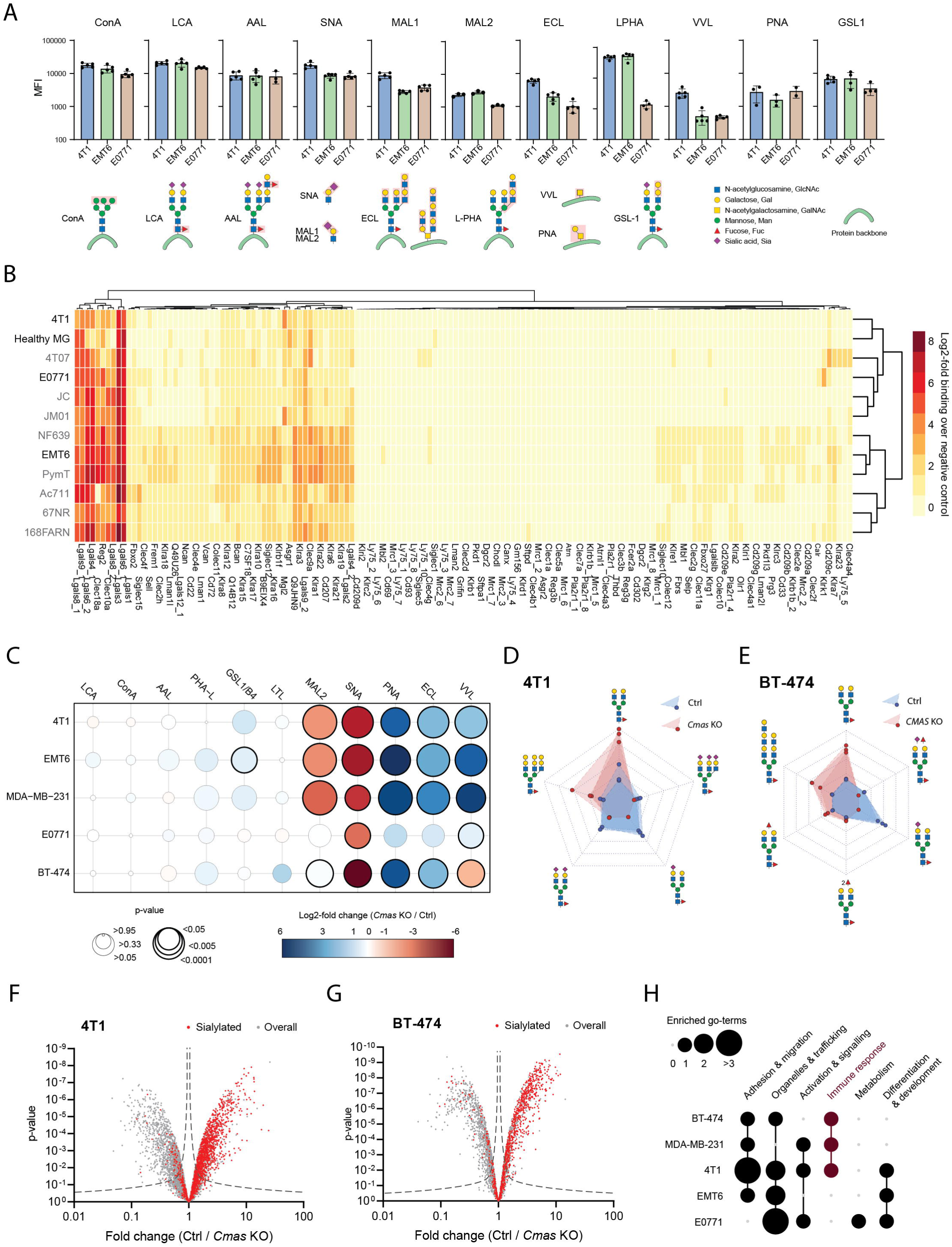
Functional and structural glycan characterization of mammary cancer cell lines. **A.** Surface glycan epitopes of the mammary cancer cells 4T1, EMT6, and E0771 assessed by cytometry using the indicated panel of 11 plant lectins. Main glycan features detected by the lectins are shown at the bottom of the panel. Results are shown as average mean fluorescence intensity (MFI) of independent replicates (n = 3-5) and their standard deviation. Concanavalin A (ConA), *Erythrina Cristagalli* Lectin (ECL), *Lens Culinaris* Agglutinin (LCA), *Sambucus Nigra* Lectin (SNA), *Maackia Amurensis* Lectin (MAL), *Vicia Villosa* Lectin (VVL), *Phaseolus Vulgaris* Leucoagglutinin (L-PHA), *Griffonia Simplicifolia* Lectin (GSL), *Aleuria Aurantia* Lectin (AAL), Peanut Agglutinin (PNA). **B.** Surface binding of 150 murine lectin-Fc fusion proteins against 11 murine mammary cancer cell lines (4T1, E0771, EMT6, 4T07, JC, JM01, NF639, PyMT, Ac711, 67NR and 168FARN) and healthy mammary epithelial cells (Healthy MG) assessed by cytometry. Results are shown as log2 fold binding over Fc control (average of 3 replicates) and detailed in Suppl. Table 1. **C.** Changes in surface glycan epitope expression in murine mammary cancer cells (4T1, EMT6, E0771) and human breast cancer cells (MDA-MB-231, BT-474) upon *CMAS* knockout. Glycan epitopes were cytometrically assessed with a panel of fluorescently labelled lectin-Fc fusion proteins. Main glycan features detected by the lectins are shown in (A). Color code reflects log2 fold change of mean fluorescence intensity and circle size negatively correlates with the p-value (n = 3, t-test). Bold circles indicate statistical significance (p < 0.05). Log2 fold changes were used for unsupervised clustering. **D and E**. Radar charts summarizing relative changes of key N-glycans comparing control and *Cmas* KO 4T1 (n = 3, D) and between control and *CMAS* KO BT-474 (n = 3, E) mammary tumor cells, as assessed by mass spectrometry. The symbol nomenclature for glycans was used to represent the glycans as detailed in (A). **F and G.** Volcano plots of glycoproteomic changes comparing control and *Cmas* knockout 4T1 (n = 3, F) and control and *CMAS* KO BT-474 (n = 3, G) mammary tumor cells. Results are shown as average fold change and p-values of the unique glycopeptides, with sialylated glycopeptides highlighted in red. Data are detailed in Suppl. Table 2. **H**. Gene ontology enrichment analysis of glycoproteins that were significantly less sialylated in *Cmas* KO cells as compared to their unedited counterparts. Sialylation status of glycoproteins was assessed via quantitative mass spectrometry (SugarQb, n = 3). Data are detailed in Suppl. Table 3.

We next conducted a screen using an extensive lectin library, covering almost the entire murine lectinome, to map the landscape of functional lectin ligands in these three breast cancer cell models (Suppl. Table 1). To our knowledge, this is the largest lectin library ever utilized in a cancer context. In addition to 4T1, EMT6, and E0771, we included healthy murine mammary epithelial cells as well as eight additional murine mammary cancer cell lines (4T07, JC, JM01, NF639, PymT, Ac711, 67NR, 168FARN) to generate a larger data set to facilitate clustering (Fig. 1B, Suppl. Table 1). Interestingly, despite the pronounced upregulation of sialylation in 4T1 cancer cells, their functional lectin interaction landscape resembled that of the normal mammary epithelium (dendrogram, Fig. 1B).

The transformation of sialic acids into the activated donor-substrate CMP-Sialic acid is mediated through the enzyme CMAS (Cytidine Monophosphate *N*-Acetylneuraminic Acid Synthetase) and is an essential step of sialylation (*9*). Subsequently, CMP-Sialic acids are transported to the Golgi apparatus, where various sialyltransferases utilize them as donor substrates to add sialic acid and modify glycans (*10, 11*). To genetically assess the role of sialylation in breast cancer, we used CRIPSR/Cas9 to knock out *Cmas* in 4T1, EMT6 and E0771 murine mammary cancer cells and in MDA-MB-231 and BT-474 human breast cancer cells. Importantly, the changes in surface glycosylation induced by *Cmas* KO were similar in mouse and human cell lines (Fig. 1C and Suppl. Fig 1B). The loss of sialylation (assessed through the loss of SNA and MAL2 binding) triggered a shift in the glycome, characterized by an increase in terminal galactose, which can be determined using the lectins ECL, PNA and GSL1 (Fig. 1C). These changes were confirmed by structural N-glycomic analysis using mass spectrometry, which shows a shift in both mouse and human cell models from terminal sialylated to terminal galactosylated N-glycans (Fig. 1D,E and Suppl. Fig. 1C,D). Subsequently, we performed comprehensive glycoproteomic analysis using our in-house developed SugarQb pipeline (*12*). This analysis enabled the quantitative assessment of 10552and 5896 unique glycopeptides, annotated to 844 and 618 glycoproteins, for mouse and human breast cancer cell lines, respectively (Fig. 1F,G; Suppl. Fig. 2A-D; Suppl. Table 2). Interestingly, the gene ontology enrichment analysis revealed that through *Cmas* KO affected glycoproteins in human breast cancer cell lines were enriched in cellular adhesion and immune response pathways, a feature reflected also by the 4T1 murine mammary cancer cell line (Fig. 1H; Suppl. Table 3). Based on these results, we focused on the 4T1 model to further assess the mechanistic impact of hyper-sialylation in mammary cancer.

### Sialylation deficiency licenses cytotoxic T cells to control mammary tumors

*In vitro* analysis of *Cmas* KO 4T1 cells and control 4T1 cells showed comparable overall growth behavior (Suppl. Fig. 3A) and morphology (data not shown). Additionally, the overall proteome remained largely unchanged (Suppl. Fig. 3B and Suppl. Table 4). However, when sialylation-deficient 4T1 cells were orthotopically implanted into the mammary fat pad of syngeneic BALB/c mice, they exhibited a significant growth disadvantage compared to control 4T1 cells (Fig. 2A). Tumor cell proliferation was similar between *Cmas* KO and control tumors, as indicated by the Ki67 proliferation marker; however, we observed a marked increase in tumor-infiltrating CD8^+^ T cells (Fig. 2B-D). Of note, *Cmas* KO 4T1 tumor cells maintained their negative sialylation phenotype *in vivo,* determined at the end of the experiment (Suppl. Fig. 3C).

**Figure 2.**
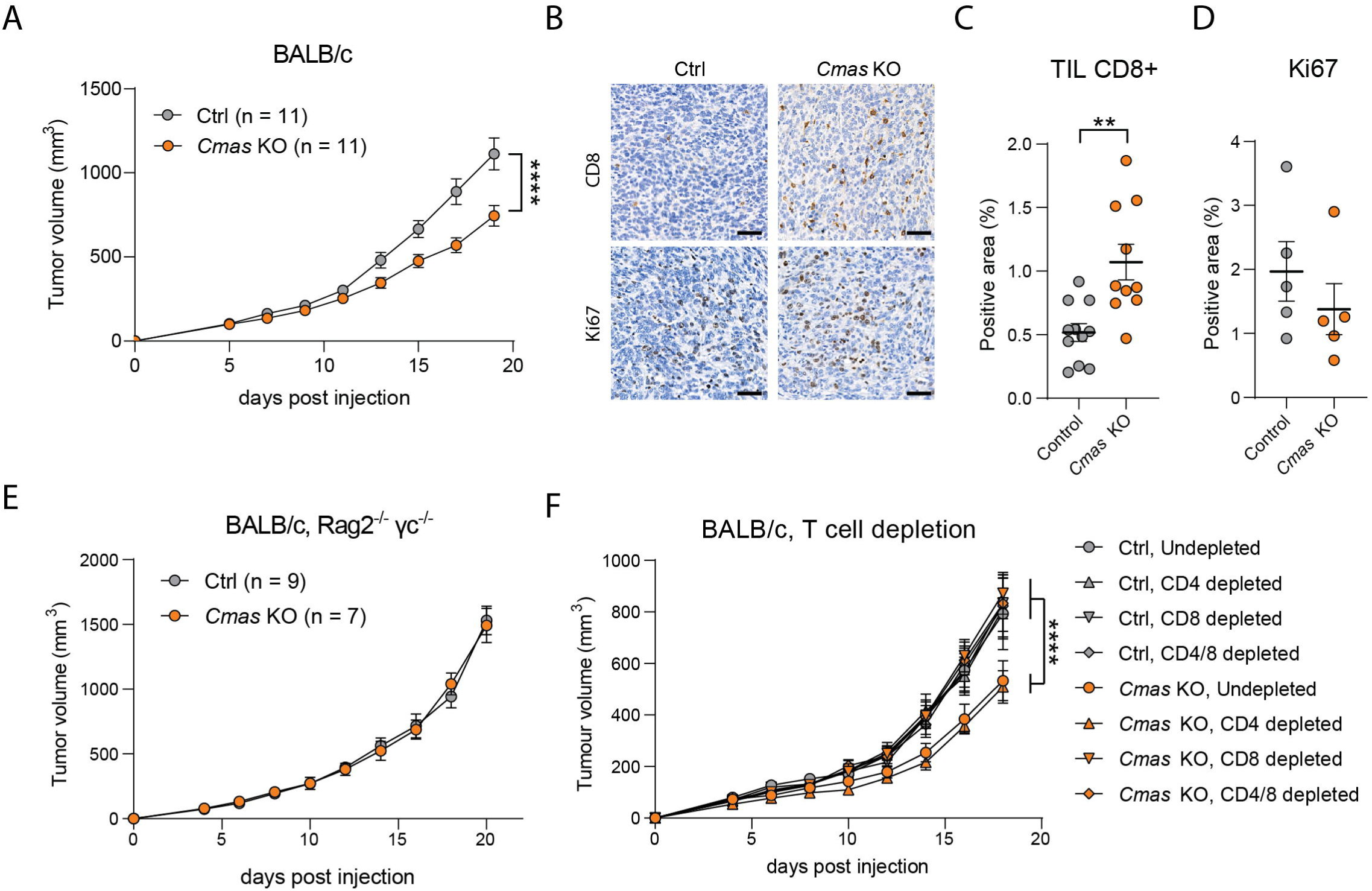
Immune-dependent *in vivo* growth differences between control and *Cmas* KO 4T1 mammary tumors. **A.** Tumor growth curves of control and *Cmas* KO 4T1 mammary tumors, orthotopically implanted into immunocompetent syngeneic BALB/c mice. **B-D**. Representative images (B) and quantification (C, D) of immunohistochemical assessment of anti-CD8 and anti-Ki67 immunostaining in Control and *Cmas* KO 4T1 mammary tumors. Scale bars correspond to 50 μm. **E**. Tumor growth curves of orthotopic control and *Cmas* KO 4T1 tumors in immunocompromised *Rag2^-/-^ γc^-/-^* BALB/c mice. **F.** Tumor growth curves of control and *Cmas* KO 4T1 tumors orthotopically implanted into syngeneic mice depleted for CD4^+^ and/or CD8^+^ T cells. n = 6 per group. Tumor growth curves are shown as mean +/-SEM and statistical differences were assessed by two-way ANOVA. Tumor volumes were determined using digital caliper. Histological quantifications are shown as mean +/- SEM and statistical difference were assessed by t-test. * p < 0.05; ** p < 0.01; *** p < 0.001, **** p < 0.0001.

To evaluate the contribution of the immune system in driving this observed phenotype, we next implanted *Cmas* KO and control 4T1 cells orthotopically into the mammary fat pad of immunodeficient *Rag2^-/-^γc^-/-^* mice, which lack B cells, T cells, and NK cells (*13*). In these immunocompromised mice, no discernible tumor cell growth differences were evident (Fig. 2E and Suppl. Fig. 3D,E). Thus, the growth disadvantage of sialylation-deficient 4T1 cancer cells stems from the host immune system. Next, to gain further insight into the role of T cells in enhancing tumor control, we depleted CD4^+^, CD8^+^, or both of these T cell populations in tumor-bearing mice (Suppl. Fig. 3F,G). Remarkably, the *in vivo* depletion of CD8^+^, but not CD4^+^ T cells restored the impaired growth of *Cmas* KO 4T1 tumors to that of control tumors (Fig. 2F). This finding establishes CD8^+^ cytotoxic T cells as a primary effector cells responsible for driving improved tumor control in sialylation-deficient 4T1 tumors.

### Loss of sialylation alters tumor infiltrating immune cell populations

To identify mechanisms that enhance tumor control in *Cmas* KO tumors, we conducted a comparative single-cell RNA sequencing (scRNASeq) analysis of tumor-infiltrating immune cells in both control and *Cmas* KO 4T1 tumors. To achieve this, we isolated CD45^+^ cells from the respective tumors, resulting in a population of more than 95% viable CD45^+^ cells (Supp Fig. 4A). The unsupervised clustering of scRNA-seq data identified 15 clusters of immune cells and one cluster of remnant cancer cells (cluster 11) (Fig. 3A, Supp Fig. 4B-E). We observed striking differences in the abundance of specific immune cell populations between control and *Cmas* 4T1 KO tumors, including a disproportionally high tumor infiltration of B cells, memory T cells, dendritic cells, and conversely a relative reduction in myeloid cells and myeloid derived suppressor cells (Fig. 3B,C). Myeloid cells (clusters 1, 4, 6, 7, and 10) of *Cmas* KO tumors remained largely unaltered in their marker expression when compared to the control tumors (Suppl. Fig. 4F). Interestingly, sub-clustering of polymorphonuclear myeloid derived suppressor cells (PMN-MDSCs, cluster 2) revealed two distinct sub-clusters, *Siglece*^+^ cluster 2A and *Ccl3*^+^ plus *Cd274*^+^ (Immune checkpoint PD-L1) cluster 2B (Fig. 3D). Particularly, the sub-cluster 2B was reduced in *Cmas* KO tumors by 23% as compared to control tumors; though these cells exhibited a comparable expression profile of the markers *Ccl3* and PD-L1 (Fig. 3D and Suppl. Fig. 4G,H). Due to low RNA content per cell, it is challenging to faithfully assess PMN-MDSCs using scRNA-seq (*14*). We therefore confirmed the significant reduction of PMN-MDSCs in *Cmas* KO tumors as compared to control tumors by flow cytometry (Fig. 3E). To assess the relevance of PMN-MDSCs accumulation in the TME, we targeted Ly6G-expressing neutrophils using depleting antibodies (Suppl. Fig. 5A-C). Importantly, the reduction in immunosuppressive PMN-MDSCs led to a significant increase in CD8^+^ T cell infiltration and a reduction in tumor growth of control 4T1 cells to a level observed in Cmas KO tumors (Fig. 3F and Suppl. Fig 5D).

**Figure 3.**
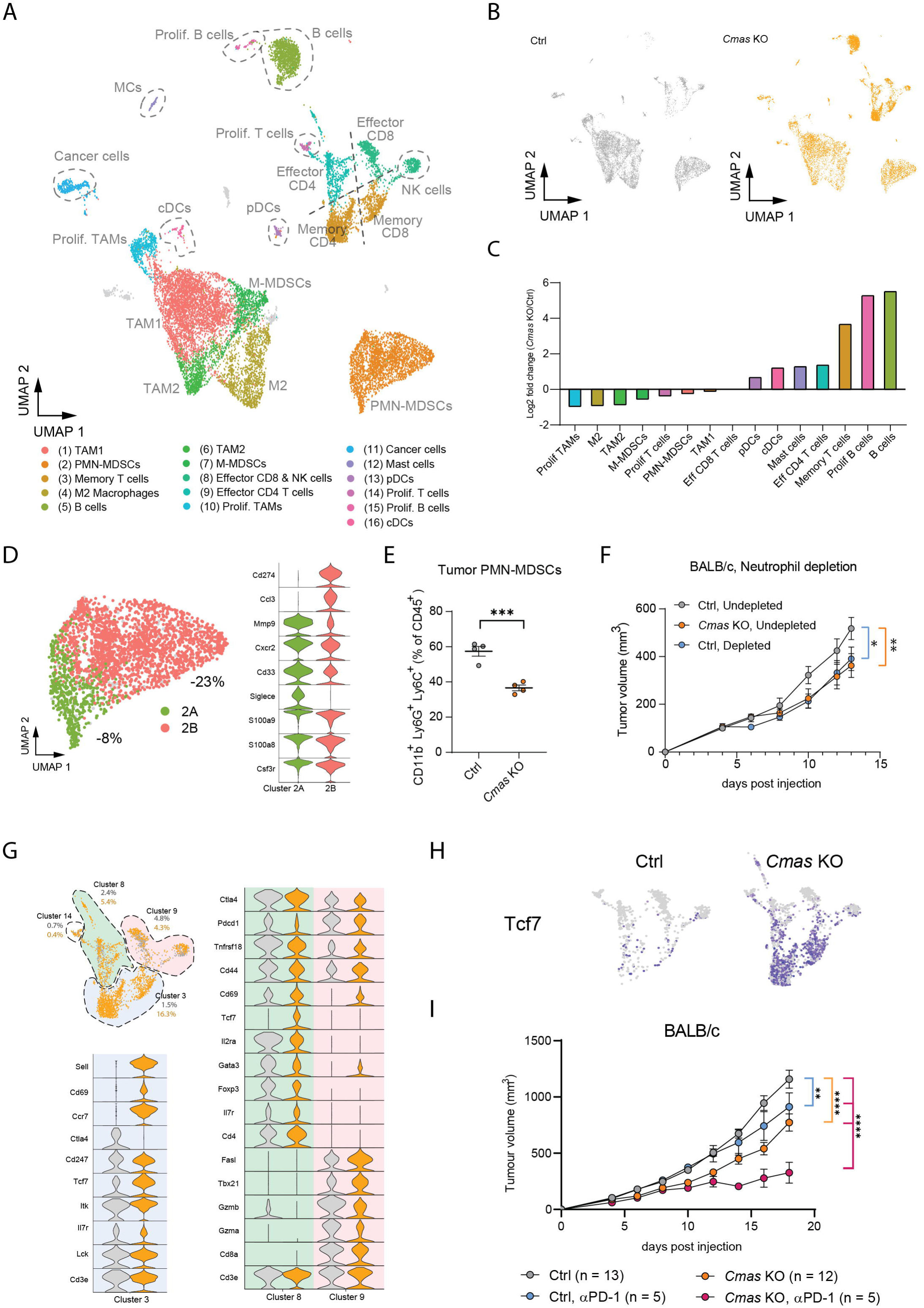
Immunologic changes in Control and *Cmas* KO 4T1 mammary tumors. **A.** Uniform manifold approximation and projection (UMAP) of the combined single cell transcriptome analyses of tumor infiltrating immune cells from control and *Cmas* KO 4T1 tumors, orthotopically implanted into immunocompetent syngeneic BALB/c mice. Clusters were annotated according to marker expression (see Suppl. Fig. 3). **B.** Individual UMAPs of tumor infiltrating immune cells from Ccontorl (top) and *Cmas* KO (bottom) 4T1 tumors. **C.** Quantitative differences in cell numbers among immune clusters comparing Control and *Cmas* KO 4T1 tumors. **D.** UMAP (left) and marker expression profiles (right) of the sub-clusters 2A (green) and 2B (red) derived from cluster 2 (polymorphonuclear myeloid derived suppressor cells, PMN-MDSC). The percentages indicate the relative reduction in cell number of respective sub-cluster from *Cmas* KO tumors as compared to control tumors. **E.** Flow cytometry-based quantification of tumor infiltrating PMN-MDSCs in control and *Cmas* KO 4T1 tumors. Results are shown as mean +/- SEM (biologic replicates, n = 4). Statistical difference was assessed by t-test. *** p < 0.001. **F.** Growth curves of control and *Cmas* KO 4T1 tumors orthotopically ionjected into BALB/c mice treated with anti-Ly6G or isotype control antibodies. n = 6-7, mean +/- SEM. Tumor volumes were determined using digital caliper. Statistical difference was assessed by two-way ANOVA (F). * p < 0.05; ** p < 0.01. **G.** Detailed UMAP of the T cell clusters 3, 8, 9 and 14 with marker expression comparisons between cells derived from control (grey) or *Cmas* KO (orange) 4T1 tumors. **H.** Feature plot of *Tcf7* expressing cells in T cell clusters 3, 8, 9 and 14 in control (left) and *Cmas* KO (right) 4T1 mammary tumors. **I.** Tumor growth of control and *Cmas* KO 4T1 tumors, orthotopically implanted into syngeneic BALB/c mice treated with anti-PD-1 or isotype control antibodies. Tumor volumes were determined using digital caliper. Histological Statistical differences were assessed by two-way ANOVA; ** p < 0.01; **** p < 0.0001.

**Figure 4.**
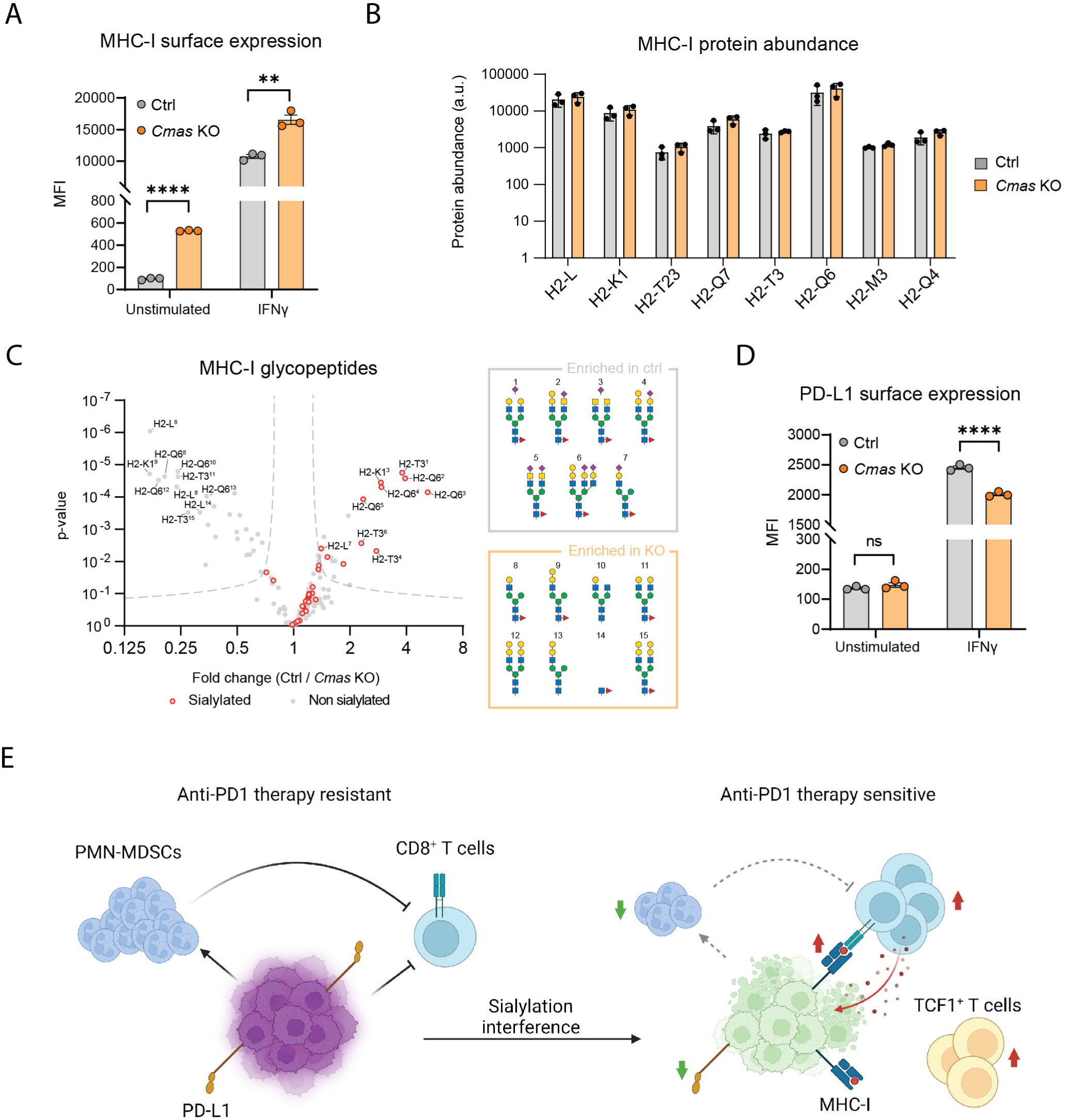
Cell autonomous changes in *Cmas* KO 4T1 cells and *in vivo* neutrophil depletion. **A.** MHC class I surface expression of unstimulated and IFNγ stimulated control and *Cmas* KO 4T1 mammary cancer cells assessed by flow cytometry detecting H-2Kd. n = 3, mean +/- SD. **B.** Quantification of the indicated MHC class I haplotype proteins in control and *Cmas* KO 4T1 cells as assessed by quantitative proteomics. n = 3, mean +/- SD. **C.** Changes in MHC class I glycopeptide abundance comparing control and *Cmas* knockout 4T1 (n = 3, F) mammary tumor cells. Results are shown as average fold change and p-values of the unique glycopeptides, with sialylated glycopeptides highlighted in red. Most significantly changed glycopeptides from the MHC-I glycoproteins H2-L, H2-Q6, H2-T3 or H2-K1 are labelled, and putative glycan structures (1–15) are annotated. Data are detailed in Suppl. Table 2. **D.** PD-L1 surface expression of unstimulated and IFNγ stimulated control and *Cmas* KO 4T1 cells assessed by anti-PD-L1 flow cytometry. n = 3, mean +/- SD. **E.** Schematic representation summarizing mechanistic alterations observed in a scenario of impaired sialylation in *Cmas* KO 4T1 tumors. MFI, mean fluorescence intensity. Statistical differences were assessed by one-way ANOVA (A, D), t-test (B) or two-way ANOVA (F). * p < 0.05; ** p < 0.01; *** p < 0.001; **** p < 0.0001.

Concomitant with the reduced number of immunosuppressive PMN-MDSCs, the numbers of defined T cell populations were markedly changed (Clusters 3, 8 and 9; Fig. 3G). In particular, *Cmas* KO tumors demonstrated a remarkable increase of *Tcf7^+^ Cd69^+^* memory T cells compared to control tumors (from 1.5% to 16.3%; Cluster 3 in blue; Fig. 3G). Of note, this cluster included both *Cd4*^+^ and *Cd8*^+^ T cells (Fig. 3A). Effector T cells, on the other hand, exhibited less pronounced changes in cell numbers while exhibiting a marked shift in their expression profiles (Cluster 8 and 9; Fig. 3G). *Cd4^+^* effector T cells (Cluster 8) in control tumors included T_H_2 cells, activated/exhausted lymphocytes, and regulatory T cells characterized by *Gata3*, *Pdcd1* and *Foxp3* expression, respectively. However, these transcription or exhaustion hallmarks were reduced in the effector T cells of *Cmas* KO tumors, which in turn expressed increased levels of the T cell activation marker *Cd69* (Cluster 8; Fig. 3G). Similarly, *Cd8^+^* effector T cells (Cluster 9) from *Cmas* KO tumors showed reduced expression of *Pdcd1* as well as increased expression of *Cd44* and *Cd69* (Fig. 3G). Importantly, among all these T cell clusters, *Tcf7* (TCF1) expression was markedly increased in *Cmas* KO tumors (Fig. 3H). In line with these findings, TCF1^+^ T cells, along with B cells, localized at the invasive tumor front of *Cmas* KO tumors, while being sparse in control tumors (Suppl. Fig. 6). In summary, abrogation of sialylation leads to reduced recruitment of PMN-MDSCs to the TME, facilitating tumor control by CD8^+^ T cells, and enables the recruitment of TCF1^+^ memory T cells to the tumor boundaries.

### Loss of sialylation sensitizes mammary tumors to anti-PD1 immunotherapy

Augmented numbers of TCF1^+^ T cells have previously been shown to correlate with increased sensitivity to immune checkpoint inhibitor therapies (*15, 16*). It is known that 4T1 tumors create a potent immunosuppressive tumor microenvironment, inhibiting CD8^+^ T cell-mediated killing and rendering anti-PD-1 immune checkpoint blockade ineffective (*17–19*). Given the enhanced anti-tumor immune response resulting from sialylation interference, in particular the accumulation of *Tcf7^+^* memory T cells, we therefore assessed the effectiveness of anti-PD-1 treatment in mice harboring control or *Cmas* KO 4T1 tumors. Remarkably, the loss of sialylation rendered 4T1 tumors sensitive to anti-PD-1 therapy, leading to prolonged tumor control and significantly improved survival in tumor-bearing mice (Fig. 3I; Suppl. Fig. 7A-D). These data show that ablation of tumor sialylation reduced PD-L1^+^ PMN-MDSC infiltration and lead to the accumulation of *Tcf7^+^* memory T cells, ultimately sensitizing resistant tumors to anti-PD-1 treatment.

### CD8^+^ T cell-mediated tumor control relies on MHC-I and PD-L1 surface expression

Next, we aimed to determine whether cell autonomous changes could possibly contribute to the CD8^+^ T cell-mediated tumor control. To assess possible alterations in the cancer cells, we measured the surface levels of key proteins involved in the interaction between cancer cells and CD8^+^ T cells, namely MHC-I and PD-L1. Interestingly, *Cmas* KO 4T1 cells showed significantly higher expression of MHC-I compared to control 4T1 cells, with or without IFNγ exposure (Fig. 4A). Notably, the total levels of different MHC haplotype proteins remained unchanged, as determined by quantitative proteomics of lysed tumor cells (Fig. 4B), indicating alterations in surface localization rather than overall expression. Consistently, the expressed glycoforms of 4 MHC haplotype proteins (H2-L, H2-K1, H2-T3, H2-Q6) changed significantly, from primarily sialylated biantennary N-glycans to terminal galactosylated and partially truncated N-glycans (Fig. 4C). In addition, the surface expression levels of MHC-I significantly increased in human breast cancer cells upon *CMAS* deletion (Suppl. Fig. 8). In contrast, cell surface expression of the immune checkpoint ligand PD-L1 on the mammary tumor cells was strikingly reduced following IFNγ stimulation of *Cmas* KO 4T1 cells (Fig. 4D). These findings indicate that alterations of tumor cell sialylation results in deregulated expression of MHC-I and PD-L1, two key factors involved in CD8^+^ T cell-mediated recognition and killing. In conjunction with the restricted accumulation of PMN-MDSCs, sialylation interference culminates in 4T1 tumors in increased infiltration of CD8^+^ T cells and better tumor control (Fig. 4E).

### Pharmacological inhibition of sialylation phenocopies *Cmas* genetic ablation for cancer immunosurveillance

Having demonstrated the potential of interfering with sialylation through genetic ablation in tumor cell lines, our next objective was to investigate whether similar effects could be achieved through pharmacologic intervention. To accomplish this, we administered 3Fax-Peracetyl N-Acetylneuraminic acid, a sialyltransferase inhibitor (STI) (*22*), to 4T1 tumor-bearing mice. *In vivo* treatment with STI resulted in a significant reduction in tumor growth (Fig. 5A). Additionally, STI licensed the tumor bearing hosts to better respond to anti-PD-1-based immunotherapy, leading to prolonged tumor control and a doubling of the overall survival (Fig. 5A and Suppl. Fig. 9A-D). Similar to genetic ablation of sialylation, pharmacological interference was dependent on an intact adaptive immune system since no spontaneous nor immunotherapy-induced tumoricidal activity was observed in *Rag2^-/-^ γc^-/-^* mice (Fig. 5B). Of note, *in vitro* experiments demonstrated that STI did not exhibit any direct cytotoxicity in 4T1 cells (Suppl. Fig. 9E).

**Figure 5.**
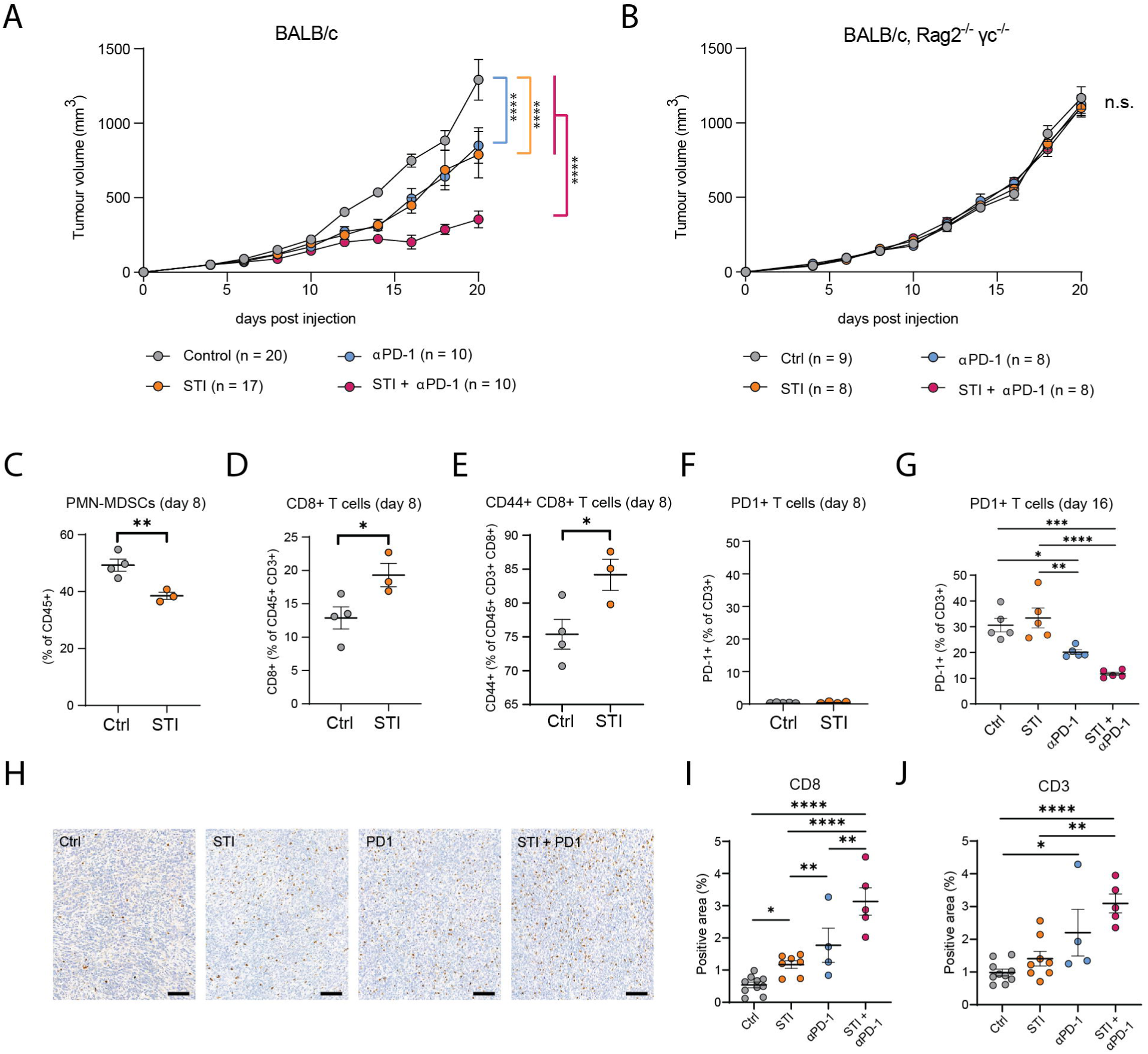
Sialyltransferase inhibitor (STI) treatment of 4T1 tumors. **A and B.** Longitudinal volumetric monitoring of orthotopic 4T1 tumor growth in immunocompetent syngeneic mice (A) or in immunocompromised *Rag2^-/-^ γc ^-/-^* mice (B) *in vivo* treated with STI and/or anti-PD-1 antibodies. Controls included treatment with vector control and/or an antibody isotype control. **C – F.** Percentages of polymorphonuclear myeloid-derived suppressor cells (PMN-MDSCs) (C), CD8^+^ T cells (D), CD44^+^ CD8^+^ T cells (E) and PD-1^+^ T cells (F) within *in vivo* orthotopic 4T1 tumors were assessed by cytometry on day 8 after tumor cell injection (n = 4, mean +/- SEM). Day 8 was chosen because at this time point there was comparable tumor growth among the STI treated and control groups. **G.** Percentages of PD-1^+^ T cells within the 4T1 tumor environment were assessed by cytometry on day 14 after initial tumor injection (n = 5, mean +/- SEM). The different treatment regimens are indicated. **H – J.** Representative images of anti-CD8 immunohistochemistry (H) and quantification (I) as well as quantification of anti-CD3 immunohistochemistry (J) within 4T1 tumors treated with STI and/or anti-PD-1 antibody. Immunohistochemistry was assessed at the endpoint when tumor volume surpassed 1500 mm^3^ (n = 4 – 10, mean +/- SEM). Scale bars = 50 μm. Statistical differences were assessed by two-way ANOVA (A, B), t-test (C - F) or one-way ANOVA (G, I, J); * p < 0.05; ** p < 0.01; *** p < 0.001; **** p < 0.0001.

Time course immune profiling revealed modulations of the tumor immune landscape upon STI treatment, consistent with those described in *Cmas* KO 4T1 tumors: STI induced a drop in PMN-MDSCs and a concomitant rise in tumor-infiltrating memory CD8^+^ T cells as determined on day 8 following tumor induction (Fig. 5C-F; Suppl. Fig. 10A). On day 14 post tumor induction, significantly fewer PD-1 expressing T cells were present in tumors treated with both STI and anti-PD-1 (Fig. 5G; Suppl. Fig. 10B). The increased infiltration of CD8^+^ T cells persisted until the endpoint of the study (Fig. 5H-J; Suppl. Fig. 10C-H). Thus, pharmacologic ablation of sialylation recapitulated the effects observed with our *Cmas* genetic ablation experiments, emphasizing the potential of sialylation interference as a therapeutic approach to license mammary tumors to immune checkpoint therapy.

### Deletion of *Cmas* in the mouse mammary gland

To further evaluate the potential of interfering with sialylation as a treatment for breast cancer, we employed an *in vivo* preclinical model of the disease. To achieve this, we established *in vivo* knockout of sialylation in the mammary gland epithelium by conditionally deleting *Cmas* using the K14-Cre delete line (*Cmas^ΔK14^*). As expected, the deletion of *Cmas* in *Cmas^ΔK14^* mice resulted in the loss of binding with sialic acid-specific lectins such as SNA, MAL1, and MAL2 (Fig. 6A and Suppl. Fig. 11A,B), confirming the generation of an effective sialylation knockout mouse line in mammary epithelial cells. Additionally, there was a significant reduction in alcian blue staining, which identifies negatively charged clusters primarily produced through sialylated mucus, when compared to control *Cmas^f/f^* mice (Fig. 6A).

**Figure 6.**
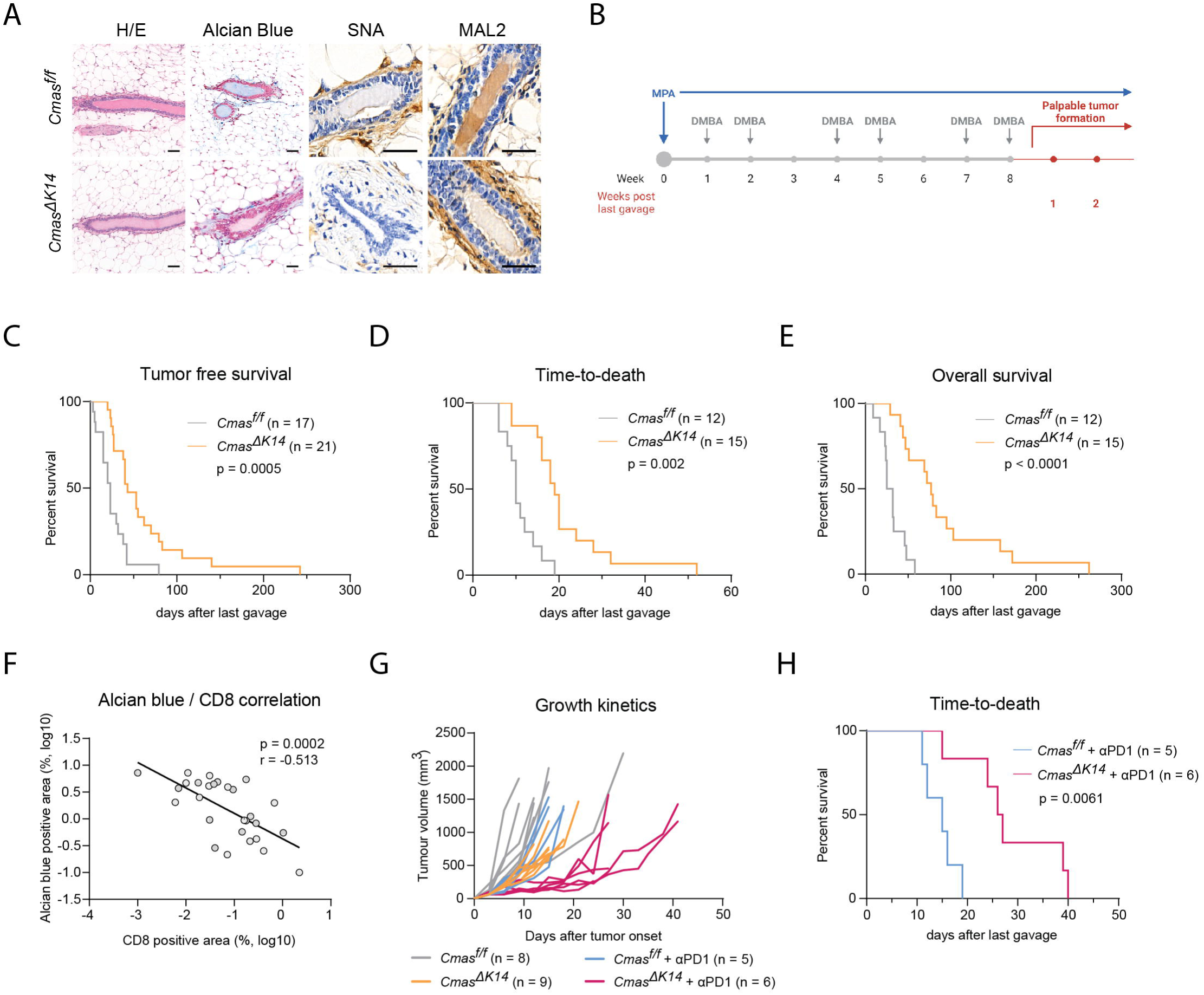
Characterization of mammary glands and spontaneous mammary tumors in *Cmas* conditional knockout mice. **A.** Histologic assessment of mammary glands stained with hematoxylin and eosin (H&E) (A), alcian blue (B), *Sambucus Nigra* lectin (SNA) (C), or *Maackia Amurensis* Lectin 2 (MAL2) (D) from nulliparous *Cmas^f/f^* and *Cmas*^Δ*K14*^ mice. Scale bars = 50 μm. **B.** Treatment scheme for the induction of spontaneous mammary tumors through the application of medroxyprogesterone acetate (MPA) slow release pellets and 7,12-dimethylbenz[a]anthracene (DMBA) oral gavages. **C – E.** Kaplan-Meier curves of tumor free survival (C), time-to-death (D), and overall survival (E) in MPA/DMBA treated *Cmas^f/f^* and *Cmas*^Δ*K14*^ mice. **F.** Correlation analysis of alcian blue and CD8 positivity in MPA/DMBA induced tumors of *Cmas^f/f^* mice. Markers were assessed by alcian blue/anti-CD8 co-staining and Spearman correlation analysis was performed to assess p and r values. **G.** Spider plot of treatment responses to anti-PD-1 or isotype control treated MPA/DMBA induced tumors in *Cmas^f/f^* and *Cmas*^Δ*K14*^ mice. **H.** Kaplan-Meier curves of time-to-death in MPA/DMBA induced and anti-PD-1 treated *Cmas^f/f^* and *Cmas*^Δ*K14*^ mice. C – D, H. p values were assessed with Log-rank (Mantel-Cox) test.

Importantly, *Cmas^ΔK14^* mice did not display any other noticeable phenotypic changes, and the development and overall histology of the mammary gland epithelium appeared normal (Fig. 6A and Suppl. Fig. 11C,D). Furthermore, the mammary glands remained functional in pregnancy and lactation, as *Cmas^ΔK14^* dams were able to nurse their pups, resulting in no reduced viability of their litter compared to control *Cmas^f/f^* dams (Suppl. Fig. 11E,F). Thus, surprisingly, deletion of *Cmas* in the mammary epithelium has no apparent effect on mammary gland development in puberty nor during lactation and pregnancy.

### Sialylation deficiency increases cancer immunosurveillance in autochthonous mammary tumors and sensitizes to PD-1 blockade

We next utilized *Cmas^ΔK14^* and control *Cmas^f/f^* mice to study the spontaneous formation of mammary tumors induced by medroxyprogesterone acetate (MPA) and 7,12-Dimethylbenz(a)anthracene (DMBA) (Fig. 6B) (*23*). The combined application of DMBA/MPA leads to the spontaneous development of aggressive mammary tumors that closely resemble many aspects of human luminal B type breast cancer, including resistance to immunotherapy (*24*). *Cmas^ΔK14^* mice, lacking sialylation in the mammary gland epithelium, exhibited a remarkable improvement in tumor-free survival (TFS) with a hazard ratio (HR) of 0.234 (95% CI: 0.103 to 0.530), time-to-death (TTD, HR of 0.126 and 95% CI: 0.0428 to 0.373), and overall survival (OS, HR of 0.085 and CI95%: 0.028 to 0.260) (Fig. 6C-E). To examine the association between sialylation and CD8^+^ T cell infiltration in *Cmas^ΔK14^* and control *Cmas^f/f^* tumors and to assess tumor architecture, we established a co-staining protocol combining alcian blue, as a surrogate marker of sialic acid (Suppl. Fig. 11G), with CD8 immunohistochemistry (Suppl. Fig. 11H). This enabled the parallel quantification of sialylation and CD8^+^ T cell infiltration in the same tumor areas. When applied to the cohort of sialylation-competent *Cmas^f/f^* mice, we observed a significant negative correlation between alcian blue positivity and CD8^+^ T cell infiltration (Fig. 6F). This finding confirms our 4T1 orthotopic tumor data, with both experimental paradigms demonstrating that high levels of sialylation act as a negative regulator of CD8^+^ T cell infiltration and tumor control in mammary tumors.

The tumors that eventually developed in *Cmas^ΔK14^* mice remained unsialylated, as evaluated by SNA and MAL2 lectin stainings, thus ruling out the possibility of “escape variants” resulting from failed Cre-recombination (Suppl. Fig 11I). This observation is consistent with our assessment using lox-stop-lox GFP reporter lines, which demonstrated a high efficiency of K14-Cre mediated recombination in tumors harboring floxed alleles (Suppl. Fig. 11J). Finally, to examine whether the loss of sialylation and the subsequent increase in tumor-infiltrating CD8^+^ T cells rendered immunotherapy-resistant mammary tumors sensitive to immunotherapy, we treated a cohort of tumor-bearing *Cmas^f/f^* and *Cmas^ΔK14^* mice with anti-PD-1. Remarkably, only *Cmas^ΔK14^* tumors exhibited a significant response to anti-PD-1 therapy as determined by reduced tumor growth (Fig. 6G) and prolonged survival (Fig. 6H, HR 0.084, CI95%: 0.014 to 0.493). These data show that genetic *in vivo* sialylation-interference apparently does not disrupt physiological mammary gland development and function, but breaks immune tolerance in immunotherapy-resistant luminal B type mammary tumors.

### Sialylation negatively correlates with CD8^+^ T cell infiltration in human breast cancer

To further validate the role of sialylation in cancer immunosurveillance and to identify breast cancer patients that might benefit from sialylation interference, we analyzed the tumor sialylation status in human breast cancer cohorts. First, we optimized histochemical staining protocols using the lectins SNA (binding α2,6 sialic acid) and MAL2 (binding α2,3 sialic acid); we consistently obtained faint positivity in the healthy human mammary gland epithelium (Suppl. Fig. 12A). The specificity of our staining protocol was confirmed using neuraminidase pretreated controls, ablating SNA and MAL2 reactivity on enzymatically desialylated sections (Suppl. Fig. 12B). Next, we assessed a cohort of 50 treatment-naïve patients diagnosed with primary breast cancer for whom pre-treatment biopsies were available (Suppl. Table 5). In this cohort, 27% and 57% of all cases showed an increased positivity for SNA or MAL2, respectively, compared to breast specimens from healthy control women. Overall, 69% of breast cancer exhibited an increase in either SNA or MAL2 specific staining, indicating a high frequency of hyper-sialylation in untreated breast cancers. We did not observe significant differences in the SNA or MAL2 staining profiles among the different molecular subtypes of breast cancer, except for Luminal A (Fig. 7A,B). Luminal A tumors consistently displayed upregulation of MAL2 staining in 8 out of 10 cases, while 7 out of 10 cases were negative for SNA staining. This unique discordant pattern suggests a selective increase of α2,3 sialylation in this molecular breast cancer subtype. Finally, we combined SNA and MAL2 staining to calculate a sialylation score for human breast cancer. No significant difference in sialylation scores was observed among the four molecular subtypes (Fig. 7C).

**Figure 7.**
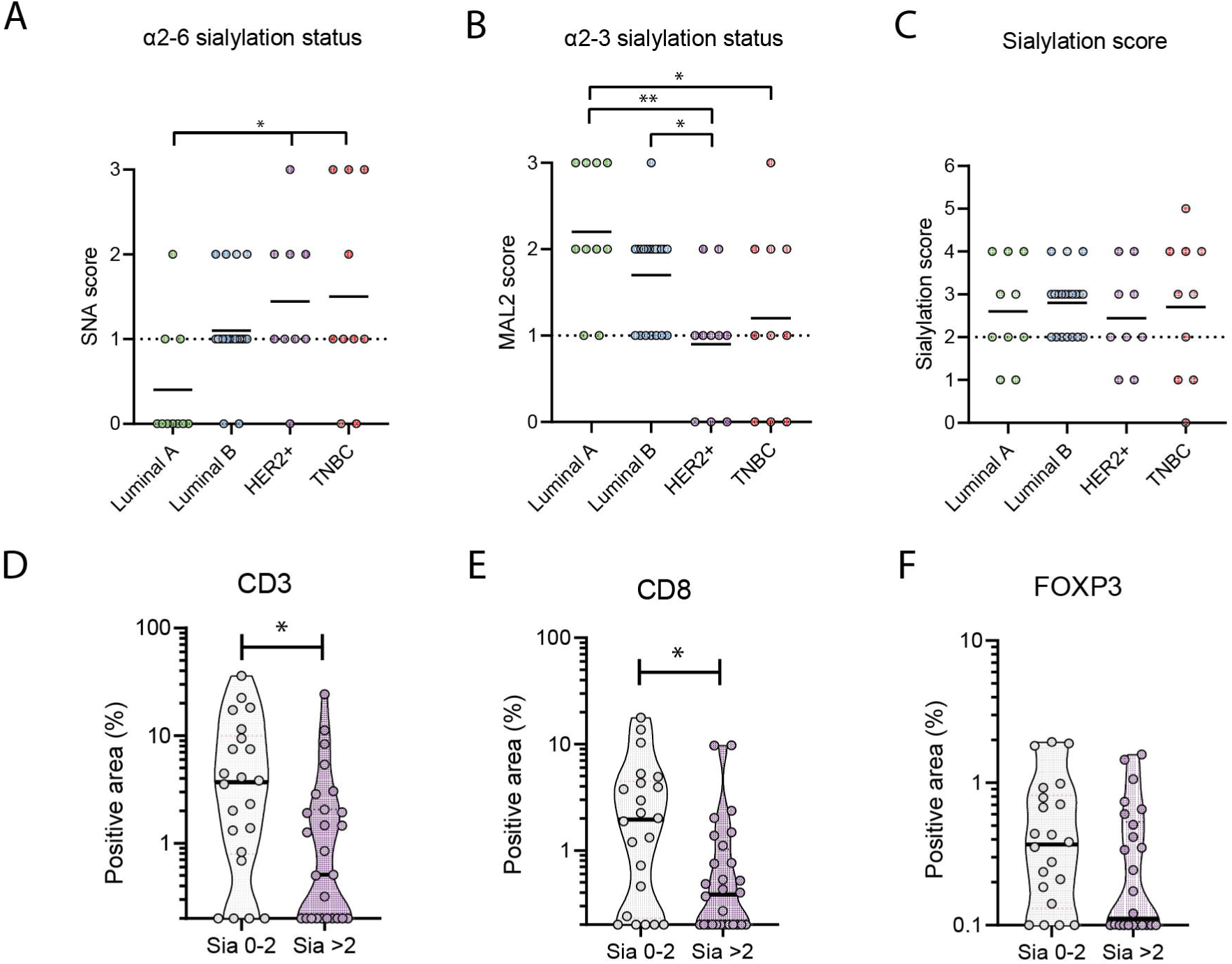
Sialylation in human breast cancer. **A and B.** Histologic evaluation of *Sambucus Nigra* lectin (SNA) (A) and *Maackia Amurensis* Lectin 2 (MAL2) (B) staining in a cohort (n = 50) of breast cancer biopsies. A score of 0, 1, 2, 3 corresponds to negative, normal amount, increased amounts, and highly increased amounts of staining, respectively, as compared to healthy breast epithelium (Suppl. Fig. 10). **C.** Sialylation score is the combined score of both SNA and MAL2 staining. A score of more than 2 corresponds to increased amounts of sialylation as compared to the healthy breast epithelium. **D – F.** Differences in CD3, CD8 and FOXP3 positive area comparing breast cancers with sialylation scores of 2 or less and scores above 2. T-test, p* < 0.05.

Lastly, we evaluated T cell infiltration, specifically using immunostaining directed against CD3, CD8, and FOXP3 (Suppl. Fig. 12C-H, Suppl. Table 5). Comparing the positivity of these T cell markers between hyper-sialylated breast tumors (sialylation score >2) and breast cancer with normal/reduced sialylation (sialylation score ≤2), a significant correlation between low levels of CD8^+^, CD3^+^ and also FOXP3^+^ T cell infiltration and hyper-sialylation was evident (Fig. 7D-F and Suppl. Fig. 12I-K). These findings in human breast cancer patients show that high tumor sialylation levels correlate with low numbers of tumor-infiltrating T cells.

## Discussion

The unique immune privilege of the breast allows marked morphological and molecular changes during development and pregnancy. Intriguingly, the immunoregulatory Siglec (Sialic acid-binding immunoglobulin-type lectins) family has expanded in mammals together with the evolution of the mammary glands (*25*). We therefore speculated that among all organs, the mammary epithelium, which is the evolutionary youngest and defining organ of mammals, may particularly capitalize on the sialic acid immunoregulatory circuits.

Surprisingly, genetic inactivation of sialylation had only minor effects on the development and function of mammary glands in mice, suggesting that sialic acids are not crucial components of physiological mammary gland biology. This is in line with previous studies that have shown that sialylation is not essential for early embryonic development and organogenesis in mice (*26, 27*). In contrast, our data indicates that the absence of sialylation has beneficial effects in tumorigenesis, as it protects against the spontaneous formation of mammary tumors upon progestin and carcinogen treatment. However, it is likely that sialylation of mammary epithelial cells plays a role in processes such as infection clearance or prevention of autoimmune diseases, which needs to be explored in future experiments. Importantly, our data indicates that the absence of sialylation has beneficial effects in tumorigenesis, as it protects against the spontaneous formation of mammary tumors upon progestin and carcinogen treatment. The lack of evident pathology in conditional knockout mice further suggests that local interference with mammary gland sialylation may be well-tolerated and non-hazardous to the non-transformed mammary tissue in breast cancer patients.

Similarly, the loss of sialylation did not affect the *in vitro* growth of any of the examined murine and human mammary tumor cell lines. This provides support for the non-essential role of sialylation in both healthy and even transformed mammary tumor cell lines. In the 4T1 mammary cancer cell line, which is widely utilized in murine mammary tumor studies, the impact of sialylation interference appears to be exclusively due to immunoregulation. This is evident from the absence of growth disparities when compared to sialylation-competent tumor cells in immunocompromised mice. Moreover, it is conceivable that the removal of sialylation also prevents metastasis, as previously described (*28*).

Mechanistically, using multiple assays such as single cell transcriptomics or immunodetection, we report that sialylation in breast tumors drives the recruitment of PMN-MDSCs while hindering the efficient eradication of tumors by CD8^+^ T cells. Consequently, pharmacologic or genetic abrogation of tumor sialylation facilitated CD8^+^mediated tumor control and the recruitment of Tcf7^+^ CD8^+^ memory T cells within the tumor microenvironment. In multiple previous studies, *Tcf7^+^ CD8^+^* memory T cells have been linked to immune checkpoint inhibitor responses (*15, 16, 29–32*). The interference with sialylation therefore resulted not only in prolonged control of the breast tumors but, most importantly, also in breaking the resistance of breast tumors to anti-PD1 therapy.

Previous studies have shown that other innate immune cells, such as tumor-associated macrophages, can contribute to improved CD8^+^ T cell-mediated tumor control upon partial desialylation of colorectal cancer and melanoma models (*6*). This phenotype was functionally linked to the actions of Siglecs (*6, 7*). In contrast, complete desialylation interference in a colorectal tumor model resulted in immune-dependent accelerated tumor growth (*33*). These observations are likely consequences of the intricate sialylation-immune axis and suggest that, similar to other immunotherapies, the outcome might depend on tumor-intrinsic properties. Thus, it is crucial to define key determinants that can serve as predictive markers for treatment outcomes following sialylation interference. For breast cancer, our data, along with findings from other studies (*6, 34, 35*), consistently demonstrate that sialylation contributes to a suppressed immune response. This is likely attributable not only to sialylation-driven recruitment and polarization of immune-dampening innate immune cells but also, as we observed in our experiments, to increased cell surface expression of MHC-I and decreased expression of PD-L1. Previous reports have shown that increased MHC-I surface retention can also be observed in desialylated dendritic cells (*36*). Collectively, sialylation in breast cancer cells appears to affect various cellular pathways and immune cells, thereby promoting immune escape of the tumor cells.

In this study, using two preclinical models of immunologically “cold” and immune checkpoint inhibitor unresponsive mammary cancers, we have demonstrated that inhibition of sialylation sensitizes tumors to combination therapy with anti-PD1 immune checkpoint inhibition. This intervention represents, to our knowledge, the first successful preclinical strategy to render aggressive MPA/DMBA mammary tumors amenable to immunotherapy. This data underscores the potential of sialylation interference strategies for future clinical combination therapies, including those currently being assessed in clinical trials (NCT05259696). Finally, the high frequency with which we observed hyper-sialylation in breast tumors of patients, along with its significant negative correlation with tumor-infiltrating T lymphocytes, suggests that a substantial number of breast cancer patients could benefit from therapeutic interventions targeting sialylation.

## Material and Methods

### Cell lines

All breast cancer (BC) cell lines were obtained from ATCC (Manassas, VA) and healthy mouse primary mammary epithelial cells were obtained from Cell Biologics (C57-6035, Chicago, IL) and cultured in DMEM High Glucose medium supplemented with 10% FCS and penicillin-streptomycin (100 U Pen/ml; 0.1 mg Strep/ml). Cells were authenticated by short tandem repeat analyses and regularly tested for mycoplasma by PCR.

### *Cmas* knockout cells

Guide RNAs against mouse or human *CMAS* were designed using the Broad Institute CRISPick tool (https://portals.broadinstitute.org/gppx/crispick/public). Reference genomes were Human GRCh38 or Mouse GRCm38, and the mechanism CRISPRko Enzyme: SpyoCas9. The top 5 crRNA spacer sequences were selected and cloned into the pX459v2 Cas9 Puro Plasmid (Addgene Plasmid #62988) using single-step golden-gate cloning. After plasmid purification, 500 ng of pX459v2:sgRNA DNA was transfected into cells using Lipofectamine2000 following the manufacturer’s instructions. The day after transfection, the media was replaced with growth medium containing 1 µg/ml Puromycin. After 48h of transient selection, the medium was replaced with standard growth medium and cells were bulk sorted on Aria III (BD, Franklin Lakes, NJ) using *Sambucus Nigra* Lectin (SNA, Vector) for sialylation negative cells (*Cmas* KO) and cells with unaltered sialylation levels (Ctrl). Three different guides were tested each to target human *CMAS* or mouse *Cmas* (Resource Table). The best performing guides assessed by the capacity of inducing SNA negative cells were human guide 1 and mouse guide 3, and were the guides used for all conditions. In every experiment, a more than 95% purity of SNA negative or positive cells was confirmed for *Cmas* KO and their respective control cells.

### Orthotopic tumor experiments

4T1 cells were orthotopically injected into the right inguinoabdominal mammary fat pad of BALB/c mice or BALB/c *Rag2^-/-^γc^-/-^* mice (both Jackson Laboratory, Bar Harbor, ME). For this purpose, cells were cultured in monolayer, harvested by trypsin, and washed with PBS. 100 μl of a 1:1 PBS and Matrigel (Corning Inc., Corning, NY) cell suspension (250,000 cells) were injected per mouse. Tumors were monitored and measured every second day starting from day 4 after injection. Tumors were measured using digital calipers; the tumor volume was calculated as length × width × height. Tumor bearing mice were sacrificed following ethical guidelines.

### *In vivo* cell depletion experiments

T cells were depleted from tumor bearing mice by the application of anti-CD4 (Clone GK1.5, in house produced) and/or anti-CD8 (Clone 2.43, in house produced) monoclonal antibodies. For this purpose, i.p. injection of 100 μg anti-CD4 and/or 100 μg anti-CD8 or a species matched (rat) isotype control antibody IgG2b (BioXCell, Lebanon, NH) were administered on days 0, 2, 8, 14 and 18 post tumor cell inoculation. On day 8 the T cell depletion was confirmed via flow cytometry using blood analysis. Neutrophils were depleted using a protocol modified from Boivin *et al*. (*37*). Briefly, tumor bearing mice were treated every other day by two i.p. injections. First 50 μg anti-Ly6g (1A8, BioXCell) or isotype control antibodies (BioXCell), followed by a second i.p. injection with 50 μg anti-Rat IgG (MAR18.5) two hours later. On day 7 the neutrophil depletion was confirmed using flow cytometry analysis of blood samples.

### *In vivo* sialyltransferase inhibition

For sialyltransferase inhibition experiments, tumor bearing mice were treated starting from day 4 post tumor inoculation. The inhibitor (3Fax-Peracetyl N-Acetylneuraminic acid (Sigma)) was administered at sub-lethal doses determined in pilot studies (*22*). For intra-tumor injections, 600 μg of 3Fax-Peracetyl N-Acetylneuraminic acid were dissolved in 30% DMSO in PBS and given every second day until day 12 post tumor induction. To avoid kidney damage, the treatment dose was lowered from day 14 post tumor induction to 300 μg of 3Fax-Peracetyl N-Acetylneuraminic acid every 4 days. The control group received vehicle control injections. For anti-PD-1 experiments, tumor bearing mice received i.p. injections of 250 μg anti-PD-1 or 250 μg isotype control in InVivoPure pH 7.0 Dilution Buffer (all BioXCell) on days 10, 12 and 14 post-tumor induction. Anti-PD-1 antibody injections were thereafter continued every 4 days.

### Flow cytometry

Tumors were resected from euthanized mice and immediately dissociated using a tumor dissociation kit and GentleMACS dissociator (both Miltenyi). Single cell suspensions were first incubated with CD16/32 Fc Block (BD) and Fixable Viability Dye eFluor^TM^ 780 (eBiosciences) for 20 min at 4°C, followed by antibody staining for 20 min at 4°C in PBS supplemented with 2% FCS using the reagents listed in the Resource Table. Antibody staining was determined using a BD Fortessa instrument (BD, Franklin Lakes, NJ) and analyzed with FlowJo (BD) and GraphPad Prism 8 (GraphPad Software, Boston, MA).

### Glycan analysis by flow cytometry

Cancer cells were harvested with 1 mM EDTA in PBS and washed 3 times in TSM buffer (20 mM Tris HCl, 150 mM NaCl, 2 mM CaCl2, 2mM MgCl2). Cells were then stained with fluorescently labelled plant lectins for 45 min at 4°C in TSM buffer using the reagents listed in the Resource Table. Staining was assessed with a BD Fortessa instrument (BD). Alternatively, cells were stained with a chimeric lectin-library as previously described (*38*) and screened on IQue Screener Plus (IntelliCyt/Sartorius, Göttingen, DE). All data was analyzed by FlowJo (BD) and GraphPad Prism 8 (GraphPad Software) or R Studio (Posit, Boston, MA).

### Human tumor tissue

Human breast cancer tissues were included from 50 patients collected at the Kantonsspital St. Gallen, Switzerland. In brief, breast cancer biopsies were taken for diagnostic purposes between September 2018 and July 2020 from patients who had consented to further scientific use of their biological samples. Ten cases were selected from each Luminal A, Luminal B, Luminal B HER2 positive, HER2 positive, and triple negative molecular subtypes, respectively. Additionally, ten control samples were chosen from patients with breast reduction surgery between August 2018 and December 2021. The study was approved by the local ethics committee (Ethics Committee of Eastern Switzerland, Project-ID 2022-00082) and conducted in adherence to the Declaration of Helsinki guidelines.

### Histologic analysis

Tumors were resected from euthanized mice and immediately fixed in neutral buffered formalin (HT5012, Sigma-Aldrich, Burlington, MA) for 48h at room temperature. Following several washes with PBS, fixed tumors were embedded in paraffin, cut at 2 μm thickness, and mounted on glass slides (Permaflex plus adhesive slides, Leica, Wetzlar, DE). Slides were deparaffinized in automatic stainer Gemini AS (Fisher Scientific, Hampton, NH) by sequential application of xylene substitute (Shandon, Thermo Scientific) and decreasing concentrations of ethanol in water. For chromogenic immunohistochemistry with antibodies, dewaxed slides were subjected to antigen retrieval with sodium citrate buffer (pH 6, see Resource Table), endogenous peroxidase inactivation with 3% H_2_O_2_ (H1009, Sigma) and were also blocked in 5% BSA in TBST. In case of lectin staining, endogenous peroxidase inactivation with 3% H_2_O_2_ was performed on dewaxed specimen followed by blocking with avidin biotin blocking solution (ab64212, Abcam) and 5% BSA in TBST. Slides were washed in TBS between steps and incubated with the respective primary antibodies or lectins (Resource Table) for 2h at room temperature. Detection system rabbit, detection system rat or streptavidin-HRP (Resource Table) were used for rabbit primary antibody, rat primary antibody or lectin staining, respectively. The chromogenic reaction was induced using the DAB substrate kit (ab64238, Abcam). Hematoxylin counter staining or hematoxylin and eosin (H&E) staining and subsequent dehydration were performed using reagents detailed in the Resource Table, using automatic stainer Gemini AS. Slides were mounted with cover slips by Tissue-TEK GLC (Sakura Finetek, Torrance, CA), air dried and scanned with Slide Scanner Pannoramic 250 (3DHistech Ltd, Budapest, HU). Whole mount staining was prepared as previously described (*39*) and stained with carmine alum and mounted using Eukitt Neo mounting medium (O. Kindler, Freiburg, DE). Alcian blue stains were performed by 30 min incubation with alcian blue (pH 2.5), followed by washing in running water and counterstained with Nuclear fast red for 3 min (Resource Table). Human sections stained with the lectins SNA or MAL2 were assessed with CaseViewer (3DHistech Ltd) and manually scored from 0 to 4 for the staining intensity of cancer cells as compared to staining intensity obtained in healthy mammary glands (Suppl. Fig. 12A). Sialylation scores were defined as the sum of SNA and MAL2 scores. Tumor sections stained with anti-CD3, anti-CD8 and anti-FOXP3 were analyzed using Fiji ImageJ. For this purpose, 3 representative intra-tumoral areas were selected per case and batch quantified using a sequence of color deconvolution (H DAB), intensity threshold, conversion to mask and area fraction measurement, shown as percentages of positively stained tumor cells.

### Single cell transcriptomics

Tumors were resected at day 15 post injection and immediately dissociated using a tumor dissociation kit and GentleMACS dissociator (both Miltenyi). CD45 positive cells were then isolated with anti-CD45 microbeads (Miltenyi) following a previously described protocol (*40*). Cells were counted using NucleoCounter NC250 (Chemometec) following the manufacturer instructions and CD45 positivity and viability measured on a BD Fortessa instrument (BD); more than 90% cells were viable and 95% of the cells were confirmed to express CD45. For each sample, one million cells were fixed for 22 hours at 4°C, quenched and stored at -80°C according to 10X genomic Fixation of Cells & Nuclei for Chromium Fixed RNA profiling (CG000478) using the Chromium Next GEM Single Cell Fixed RNA Sample preparation kit (PN-1000414, 10X Genomics, Pleasanton, CA). 250,000 cells per sample were used for probe hybridization using the Chromium Fixed RNA Kit, Mouse Transcriptome, 4rxn x 4BC (PN-1000496, 10X Genomics), pooled at equal numbers and washed following the Pooled Wash Workflow following the Chromium Fixed RNA Profiling Reagent kit protocol (CG000527, 10X Genomics). GEMs were generated on Chromium X (10X Genomics) with a target of 10,000 cells recovered and libraries prepared according to the manufacturer instructions (CG000527, 10X Genomics). Sequencing was performed using NovaSeq S4 lane PE150 (Illumina) with a target of 15,000 reads per cell. Alignment of the samples was perfomed with the 10x Genomics Cell Ranger 7.1.0 multi pipeline (*41*). The reads were trimmed to a length of 28bp read1 and 92bp read2 and aligned to the mouse reference genome (refdata-gex-mm10-2020-A) and probeset (Probe_Set_v1.0.1_mm10-2020-A.csv) with default parameters using cellranger multi. The four per sample count outputs were combined with cellranger aggr. The cellranger output files for each sample (filtered barcodes, features and count matrix files) were then used for custom analysis in R using the Seurat package (version 4.2.0) (*42, 43*). In brief, Seurat objects for each file were generated by keeping cells with more than 200 features. After data normalization (NormalizeData, default setting) and scaling (ScaleData, default settings), the top 2000 variable genes were computed using FindVariableFeaturesgenes (“vst” method). For each sample, data was then transformed using SCTransform (SCTransform, vst.flavor = “v2“), followed by principal component analysis (RunPCA, npcs=50). Data integration was then performed by first generating a list containing the individual sample object, followed by selection of the top 3000 most variable features (SelectIntegrationFeatures, nfeatures = 3000) and final integration using PrepSCTIntegration (anchor.features = features from SelectIntegrationFeatures). FindIntegrationAnchors (normalization.method = “SCT“) was used to find combined integration anchors, which were then used to performa final integration using IntegrateData (normalization.method = “SCT“). RunPCA was used to find the top principal compoments of the integrated data object for downstream clustering. The first 40 PCAs were used to find neighbours FindNeighbors (reduction = “pca”, dims = 1:40), followed by clustering (FindClusters, resolution=0.7 or 0.5 or 0.3) and UMAP dimensionalty reduction (RunUMAP reduction = “pca”, dims = 1:40). To assign clusters to cell types, FindAllMarkers was used to find features that are upregulated in individual clusters with a log-Fold Change of at least 0.25, and which are expressed in at least 25% of cells (logfc.threshold = 0.25,min.pct = 0.25). The list was then used to manually assign cell types. R-code, raw sequencing files and Seurat object are available via (NCBI GEO).

### N-Glycomic analyses

Cancer cells were harvested using 1mM EDTA in PBS and cell pellets were washed once with TSM buffer (20 mM Tris HCl, 150 mM NaCl, 2 mM CaCl2, 2mM MgCl2) and further three times with PBS. Cell pellets were then mixed with 100 mM ammonium bicarbonate buffer containing 20 mM dithiothreitol and 2 % sodium dodecyl sulphate at a total volume of 500 µL. After homogenization with an Ultra-Turrax T25 disperser, the homogenates were incubated at 56°C for 30 min. After cooling down, the solution was brought to 40 mM iodoacetamide and incubated at room temperature in the dark for 30 min. Chloroform-methanol extraction of the proteins was carried out using standard protocols (*44*). In brief, supernatants were mixed with four volumes of methanol, one volume of chloroform and three volumes of water in the given order. After centrifugation of the mixture for 3 min at 2500 rpm, the upper phase was removed, 4 mL of methanol were added, and pellets resuspended. The solution was centrifuged, the supernatant removed, and the pellet was again resuspended in 4 mL methanol. The last step was repeated two more times. After the last methanol washing step, the pellet was dried at room temperature. Dried pellets were resuspended in 50 mM ammonium acetate (pH 8.4). For N-glycan release 2.5 U N-glycosidase F (Roche, DE) were added and the resulting mixture was incubated overnight at 37°C. The reaction mixture was acidified with some drops of glacial acetic acid and centrifuged at 4000×g. The supernatant was loaded onto a Hypersep C18 cartridge (1000 mg; Thermo Scientific, Vienna) that had been primed with 2 mL of methanol and equilibrated with 10 mL water. The sample was applied, and the column was washed with 4 mL water. The flow-through and the wash solution were collected and subjected to centrifugal evaporation. Reduction of the glycans was carried out in 1 % sodium borohydride in 50 mM NaOH at room temperature overnight. The reaction was quenched by the addition of two drops glacial acetic acid. Desalting was performed using HyperSep Hypercarb solid-phase extraction cartridges (25 mg) (Thermo Scientific, Vienna). The cartridges were primed with 450 μL methanol followed by 450 μL 80% acetonitrile in 10 mM ammonium bicarbonate. Equilibration was carried out by the addition of three times 450 μL water. The sample was applied and washed three times with 450 μL water. N-glycans were eluted by the addition of two times 450 μL 80% acetonitrile in 10 mM ammonium bicarbonate. The eluate was subjected to centrifugal evaporation and the dried N-glycans were resuspended in 20 μL HQ-water. 5 μL of each sample were subjected to LC-MS/MS analysis. LC-MS analysis was performed on a Dionex Ultimate 3000 UHPLC system coupled to an Orbitrap Exploris 480 Mass Spectrometer (Thermo Scientific). The purified glycans were loaded on a Hypercarb column (100 mm × 0.32 mm, 5 μm particle size, Thermo Scientific, Waltham, MA, USA) with 80 mM ammonium formate as the aqueous solvent A and 80 % acetonitrile in 80 mM ammonium formate as solvent B. The gradient was as follows: 0 – 4.5 min 1 % solvent B, from 4.5 – 5.5 min 1 – 9 % B, from 5.5 – 30 min 9 - 20 % B, from 30 – 41.5 min 20 - 35 % B, from 41.5 – 45 min 35 – 65 % B, followed by an equilibration period at 1 % solvent B from 45–55 min. The flow rate was 6 μL/min. MS analysis was performed in data-dependent acquisition (DDA) mode with positive polarity from 500 – 1500 m/z using the following parameters: resolution was set to 120,000 with a normalized AGC target of 300%. The 10 most abundant precursors (charge states 2-6) within an isolation window of 1.4 m/z were selected for fragmentation. Dynamic exclusion was set at 20 s (n=1) with a mass tolerance of ± 10 ppm. Normalized collision energy (NCE) for HCD was set to 20, 25 and 30% and the precursor intensity threshold was set to 8000. MS/MS spectra were recorded with a resolution of 15,000, using a normalized AGC target of 100% and a maximum accumulation time of 100 ms.

### Cell lysis for glycoproteomics analysis

All cell pellets were lysed using a buffer containing 50 mM Tris-HCl (Trizma Base: Sigma-Aldrich, Cat#T6066; HCl: Sigma-Aldrich, Cat#258148), 8M urea (VWR, Cat#0568), 150 mM NaCl (Merck, Cat#106404), pH 8, containing phosphatase inhibitors (Cell Signaling Technology, Cat#5870). Samples were kept on ice for 30 minbefore being sonicated using a water bath sonicator for 1 min at low intensity. Samples were centrifuged at 20,000 x g for 15 min at 4°C. Resulting supernatants were transferred to a new tube, and protein contents were quantified using Pierce™ BCA Protein Assay Kit (Thermo Fischer, Cat#23225). For each sample, 400 µg of protein were reduced using DTT (Roche, Cat#10708984001; final concentration 10 mM) for 30 min at 37°C, and subsequently alkylated using IAA (Sigma-Aldrich, Cat#I6125; final concentration 20 mM) for 30 min in the dark at room temperature. Samples were then diluted using a 50 mM Tris-HCl buffer to lower urea concentrations to 4M before adding 8 µg of Lys C (FUJIFILM Wako, Cat#125 05061) to each sample (1 µg Lys C: 50 µg sample protein). Samples were incubated for 2 h at 25°C, before being further diluted by adding 50 mM Tris-HCl buffer to lower the urea concentration to 2M. 8 µg of Trypsin Gold (Promega, Cat#V5280) were added to each sample (1 µg Trypsin: 50 µg sample protein), and samples were further incubated for 15 h at 32°C. At this point, samples were acidified using TFA to a final concentration of 0.3% before being cleaned using 200 mg (3cc) Sep-Pak cartridges (Waters, Cat#WAT054945). The resulting elution fraction containing the cleaned peptides was dried under vacuum and stored at 20°C before proceeding with the TMT labelling protocol.

### TMT labelling before glycoproteomics analyses

After quantification, 120 µg of peptides of each sample were prepared in a final volume of 100 µL of 100 mM HEPES buffer pH 7.6. Samples were labeled using TMTpro 18 plex (Thermo Scientific, Cat#A52045) reagents, according to manufacturer’s instructions. The labelling efficiency was determined by LC-MS/MS on a small aliquot of each sample. After quenching the labeling reaction, samples were mixed in equimolar amounts, evaluated by LC-MS/MS. The mixed sample was acidified to a pH below 2 with 10% TFA and desalted using C18 cartridges (Sep-Pak Vac 1cc (200mg), Waters, Cat#WAT054945). Peptides were eluted with 2 x 600 μl 80% Acetonitrile (ACN) and 0.1% Formic Acid (FA), followed by freeze-drying.

### Glycopeptide enrichment, LC-MS/MS and data analyses

Glycopeptides from the resulting TMTpro mixes were enriched by performing off-line ion pairing (IP) HILIC chromatography, as described previously (*12*). Briefly, the dried samples were taken up in 100 μL 75% acetonitrile containing 0.1 % TFA, and subjected to chromatographic separation on a TSKgel Amide-80 column (4.6 x 250 mm, particle size 5μ) using a linear gradient from 0.1% TFA in 80% acetonitrile to 0.1% TFA in 40% acetonitrile over 35 min (Dionex Ultimate 3000, Thermo). The 25 collected fractions were vacuum dried. Samples were resuspended using 0.1% trifluoroacetic acid (TFA, Thermo Scientific, Cat#28903). The IP-HILIC fractions were individually analysed by LC–MS/MS. The nano HPLC system used was an UltiMate 3000 HPLC RSLC nano system (Thermo Scientific) coupled to a Q Exactive HF-X mass spectrometer (Thermo Scientific), equipped with a Proxeon nanospray source (Thermo Scientific). Peptides were loaded onto a trap column (Thermo Scientific, PepMap C18, 5 mm × 300 μm ID, 5 μm particles, 100 Å pore size) at a flow rate of 25 μL/min using 0.1% TFA as mobile phase. After 10 min, the trap column was switched in line with an analytical column (Thermo Scientific, PepMap C18, 500 mm × 75 μm ID, 2 μm, 100 Å). Peptides were eluted using a flow rate of 230 nl/min and a binary 180 min gradient. The two steps gradient started with the mobile phases: 98% A solution (water/formic acid, 99.9/0.1, v/v) and 2% B solution (water/acetonitrile/formic acid, 19.92/80/0.08, v/v/v), which was then increased to 35% B over the next 180 min, followed by a gradient of 90% B for 5 min, which was finally, in a 2 min period, decreased to the gradient 95% A and 2% B for equilibration at 30°C.

The following parameters were used for MS acquisition using an Exploris 480 instrument, operated in data-dependent mode: the instrument was operated in a positive mode; compensation voltages (CVs) used = CV-40, CV-50, CV-60; cycle time (per CV) = 1 sec. The monoisotopic precursor selection (MIPS) mode was set to Peptide. Precursor isotopes and single charge state precursors were excluded. MS1 resolution1=160,000; MS1 Normalised AGC target1=1300%; MS1 maximum inject time1=1501msec. MS1 restrictions were relaxed when few precursors were present. MS1 scan range1= 400–1,6001m/z; MS2 resolution1=145,000; MS2 Normalised AGC target1=1200%. Maximum inject time1=12501msec. Precursor ions charge states allowed = 2-7; isolation window1=11.41m/z; fixed first mass1=11101m/z; HCD collision energies1=130,33,36; exclude isotopes1=1True; dynamic exclusion1=1451s. All MS/MS data were processed and analysed using Xcalibur v3.1 (Thermo), and Byonic (Protein Metrics) included in Proteome Discoverer v3.0. Two default glycan databases (Mammalian O-glycans and Human N-glycans) were used together with the latest human or mouse UniProt database, to generate the glycopeptide search space. We used an in-house developed R script to filter out poorly matching spectra, considering a Byonic score higher than 200. For gene ontology enrichment analysis the GOrilla tool was applied (https://cbl-gorilla.cs.technion.ac.il/), which allows for the comparison of a target gene list against a background gene list. The target gene list consisted of significantly (p < 0.05) and more than two-fold changed glycopeptides (control vs *Cmas* KO) and the background gene list of all glycopeptides identified (Suppl Table 2 and 3).

### Statistical analyses

Statistical analysis was performed using GraphPad Prism 8 software (VERSION 8.1.1, GraphPad Software Inc.). The statistical comparison between two or more groups was performed using Student’s t-test or one-way ANOVA, respectively. For the comparison of longitudinal tumor growth two-way ANOVA was applied. Survival differences and hazard ratios were assessed using Kaplan–Meier method with log-rank (Mantel-Cox) test and Mantel-Haenszel method. Data consisting of technical or biological replicates are expressed as mean1±1standard deviation (SD) or mean ±1standard error of the mean (SEM), respectively, unless specified otherwise. Clinical data are expressed as violin plots or by the median. In addition, wherever possible individual values are plotted. Differences between groups were considered significant when p1<10.05.

## Supporting information

Supplementary Tables 1-4 and Resource Table

## Acknowledgments

We thank all members of our laboratories for their support and help. We thank all in-house core facilities, especially BioOptics Facility, Proteomics Facility, Molecular Biology Service, as well as the VBCF Histology facility and Next Generation Sequencing facility. SM received funding from the European Union’s Horizon 2020 research and innovation programme under the Marie Sklodowska-Curie grant agreement No 841319 and the ESPRIT-Programme of the Austrian Science Fund (FWF, Project number: ESP 166). GJ is supported by a DOC fellowship from the Austrian Academy of Sciences. OHA received funding from the Swiss National Science Foundation (SNSF, grant No P400PM_194473) and the Bruno Bloch Foundation. Work in JMP’s laboratory is supported by the Austrian Federal Ministry of Education, Science and Research, the Austrian Academy of Sciences, the City of Vienna and grants from the European Research Council (ERC Advanced grant 341036), the Austrian Science Fund (FWF Wittgenstein award Z 271-B19), the T. von Zastrow foundation, and a Canada 150 Research Chairs Program (F18-01336).

## Declaration of generative AI and AI-assisted technologies in the writing process

During the preparation of this work the authors used Chat GPT 3.5 in order to improve the grammar of the initial manuscript draft. After using this tool, the authors reviewed and edited the content and take full responsibility for the content of the publication.

