## Supplementary figures and images for "Tumor sialylation controls effective anti-cancer immunity in breast cancer"

### S Figure 1.jpg

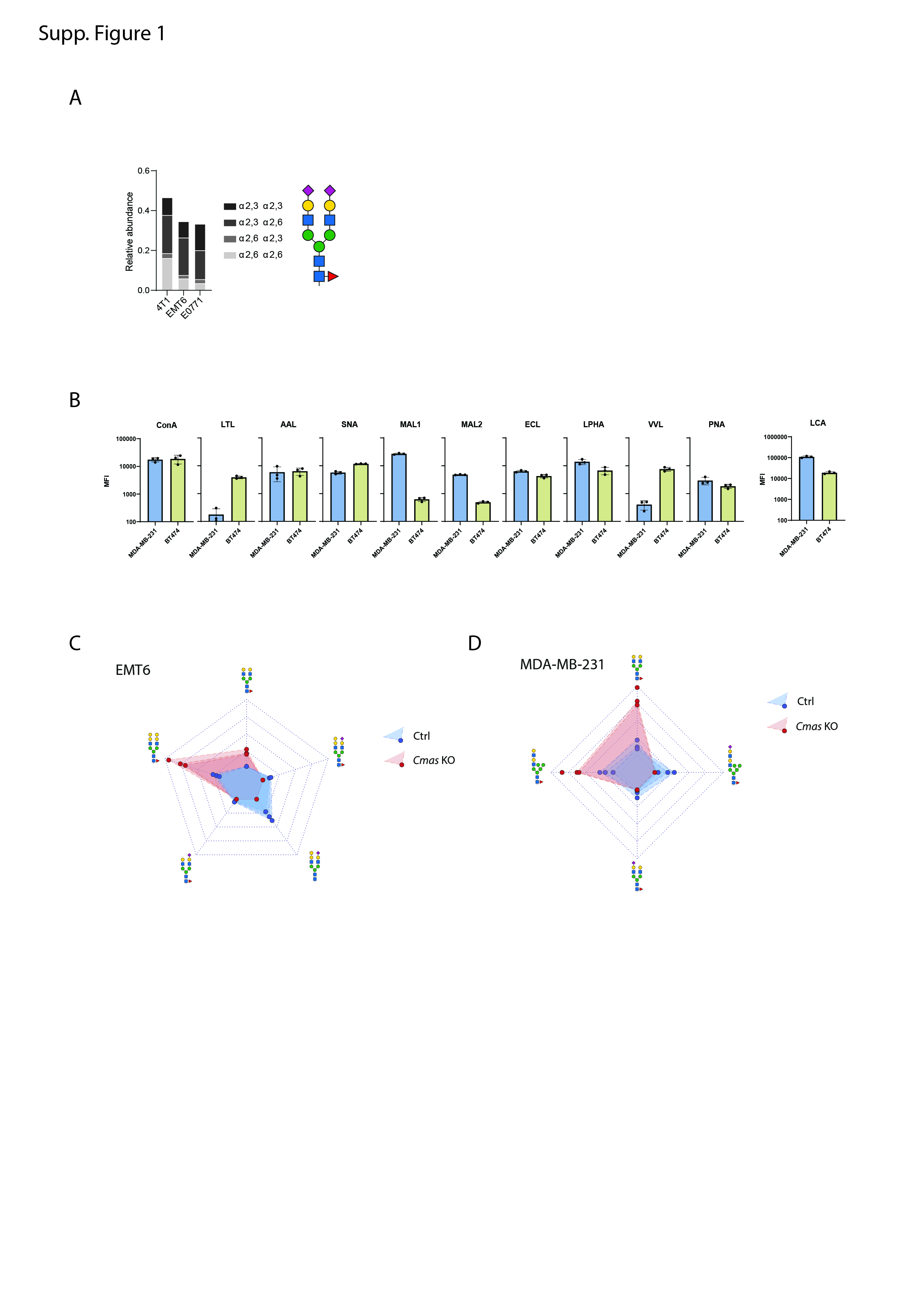

### S Figure 2.jpg

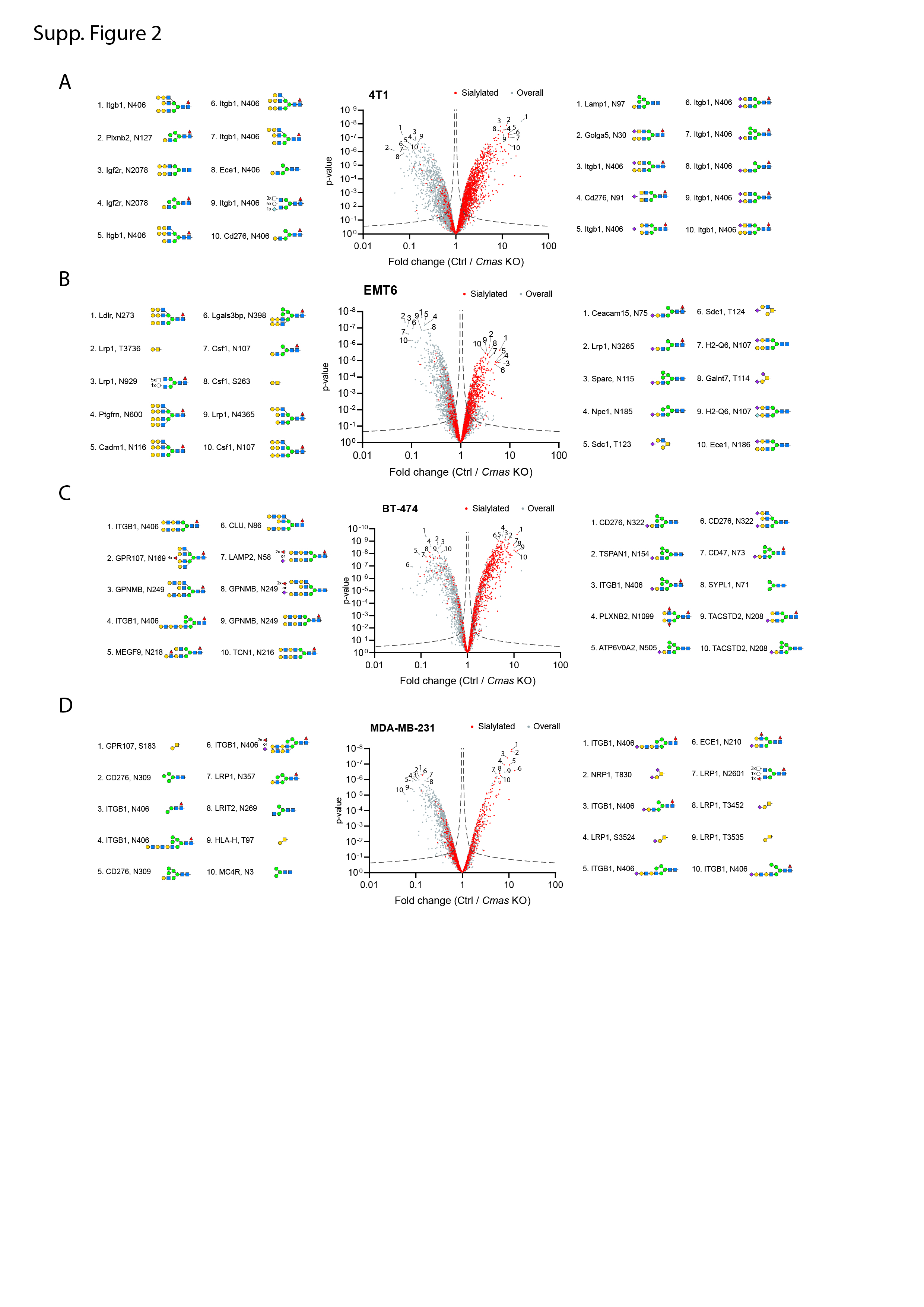

### S Figure 3.jpg

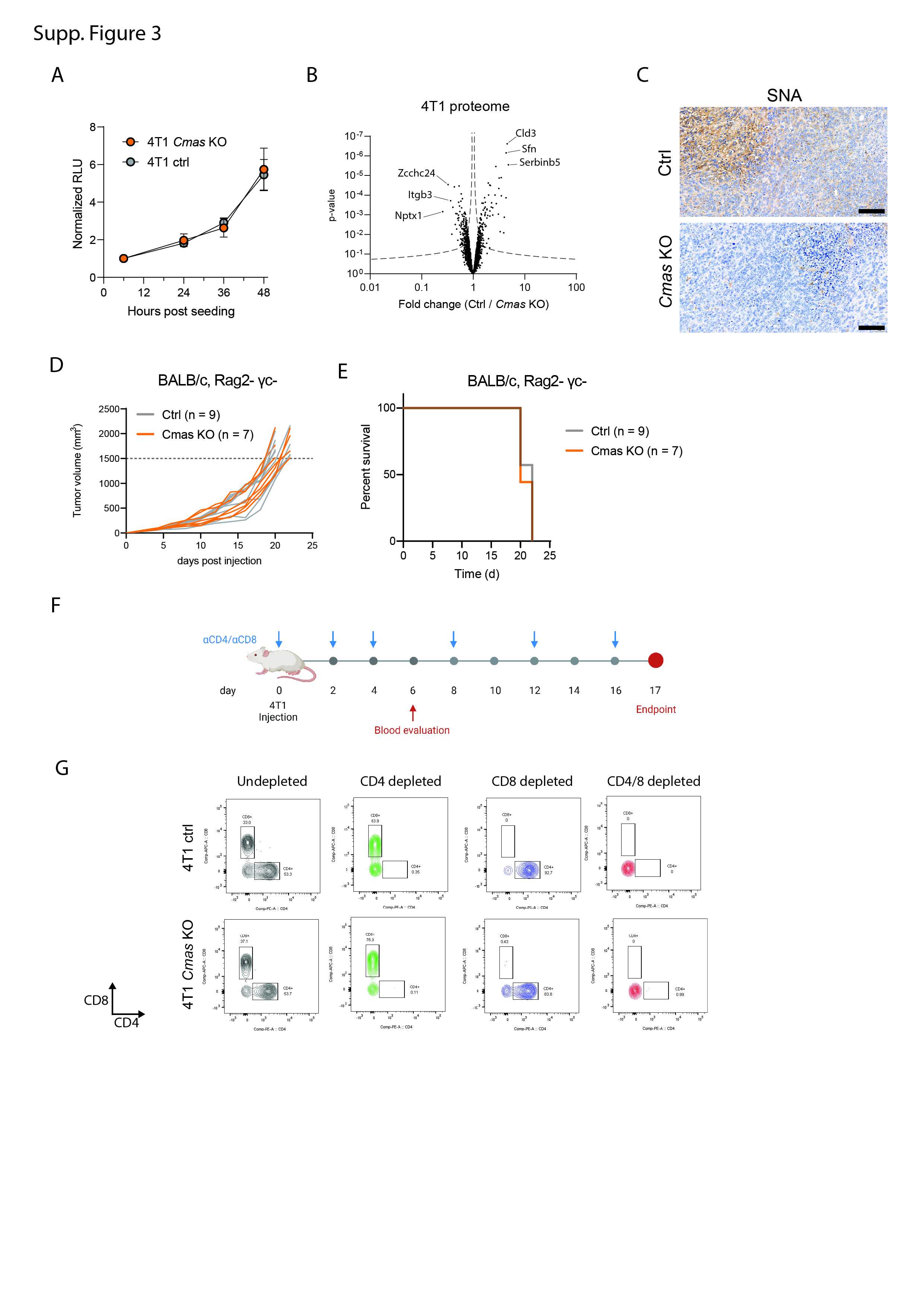

### S Figure 4.jpg

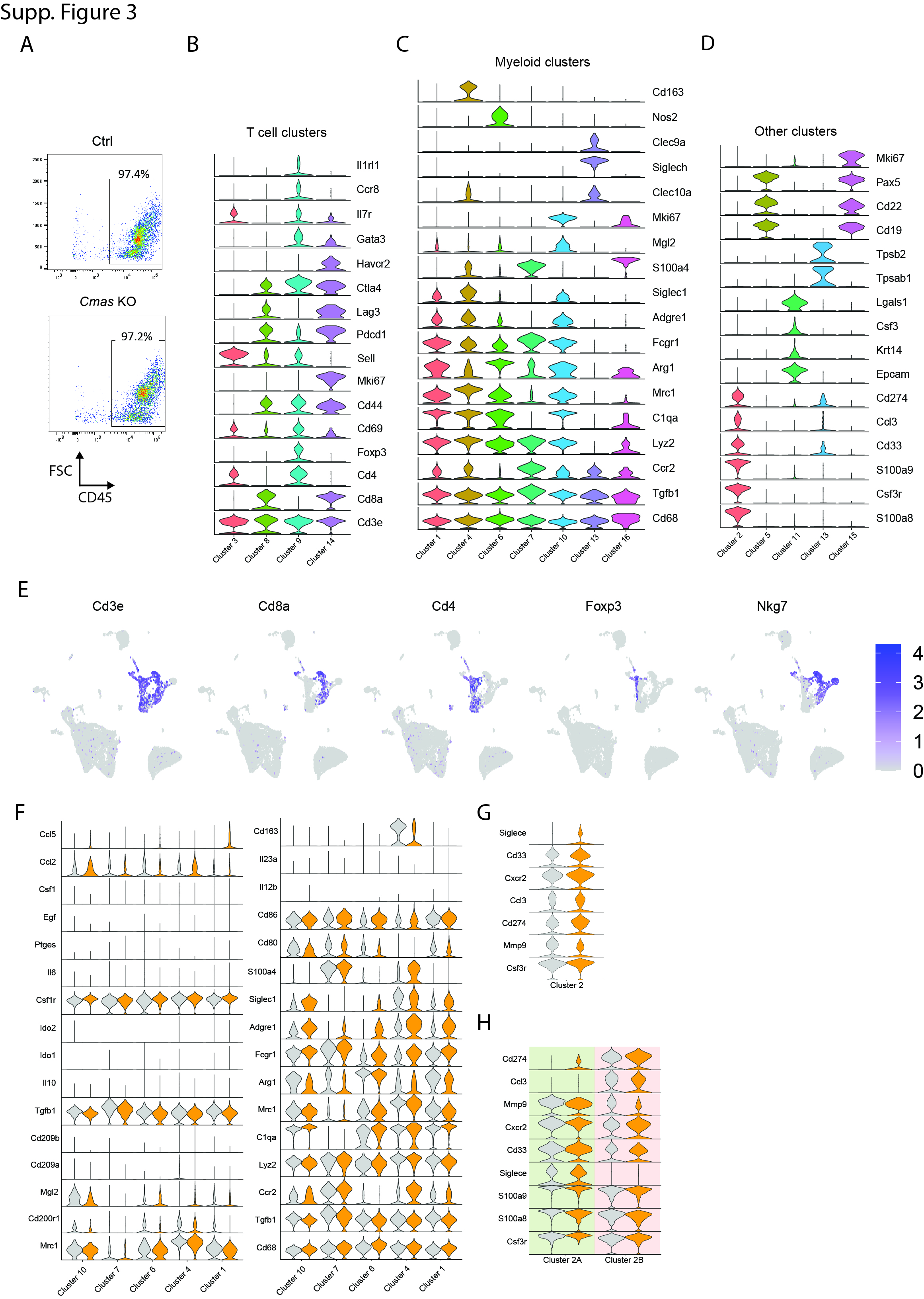

### S Figure 5.jpg

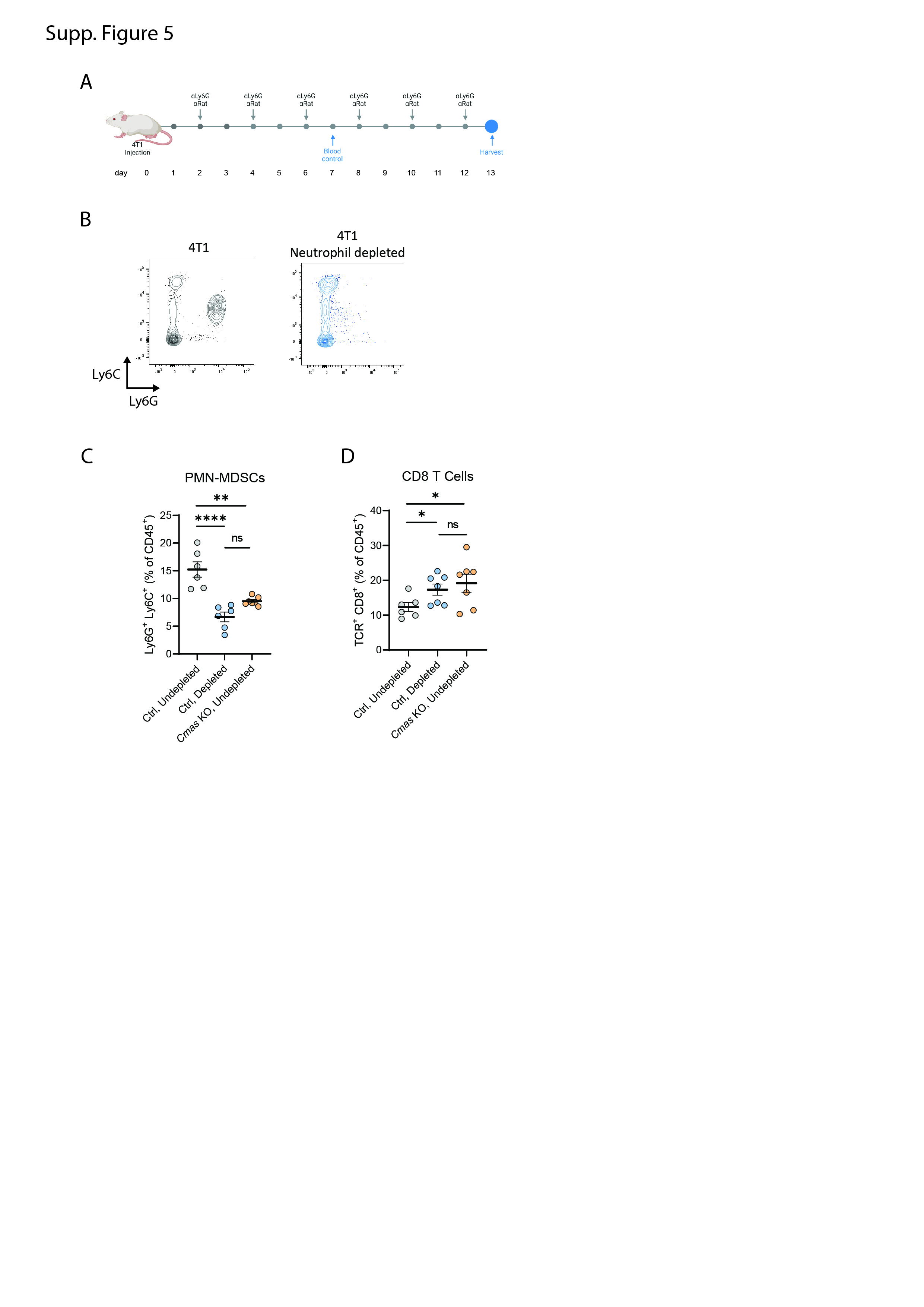

### S Figure 6.jpg

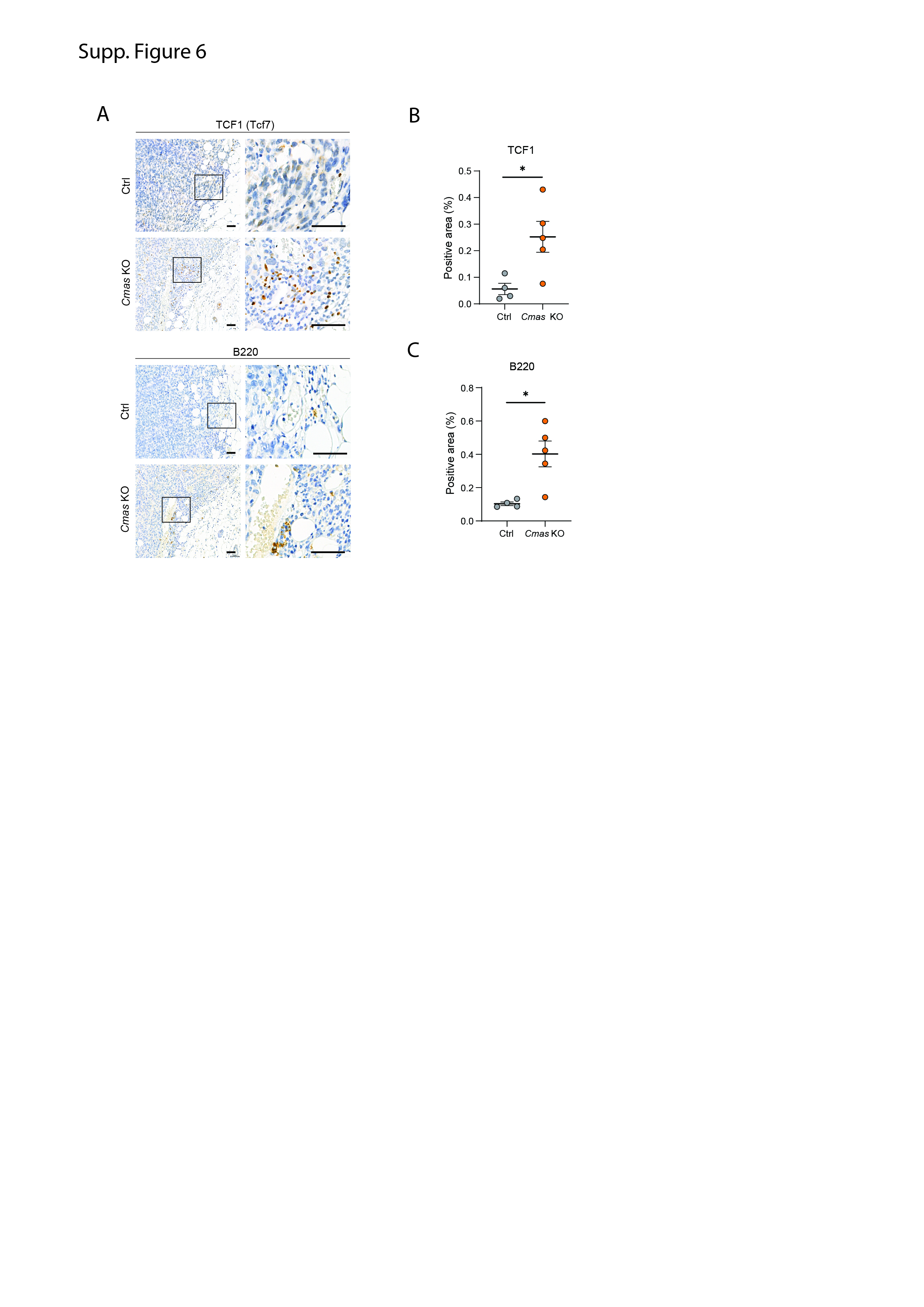

### S Figure 7.jpg

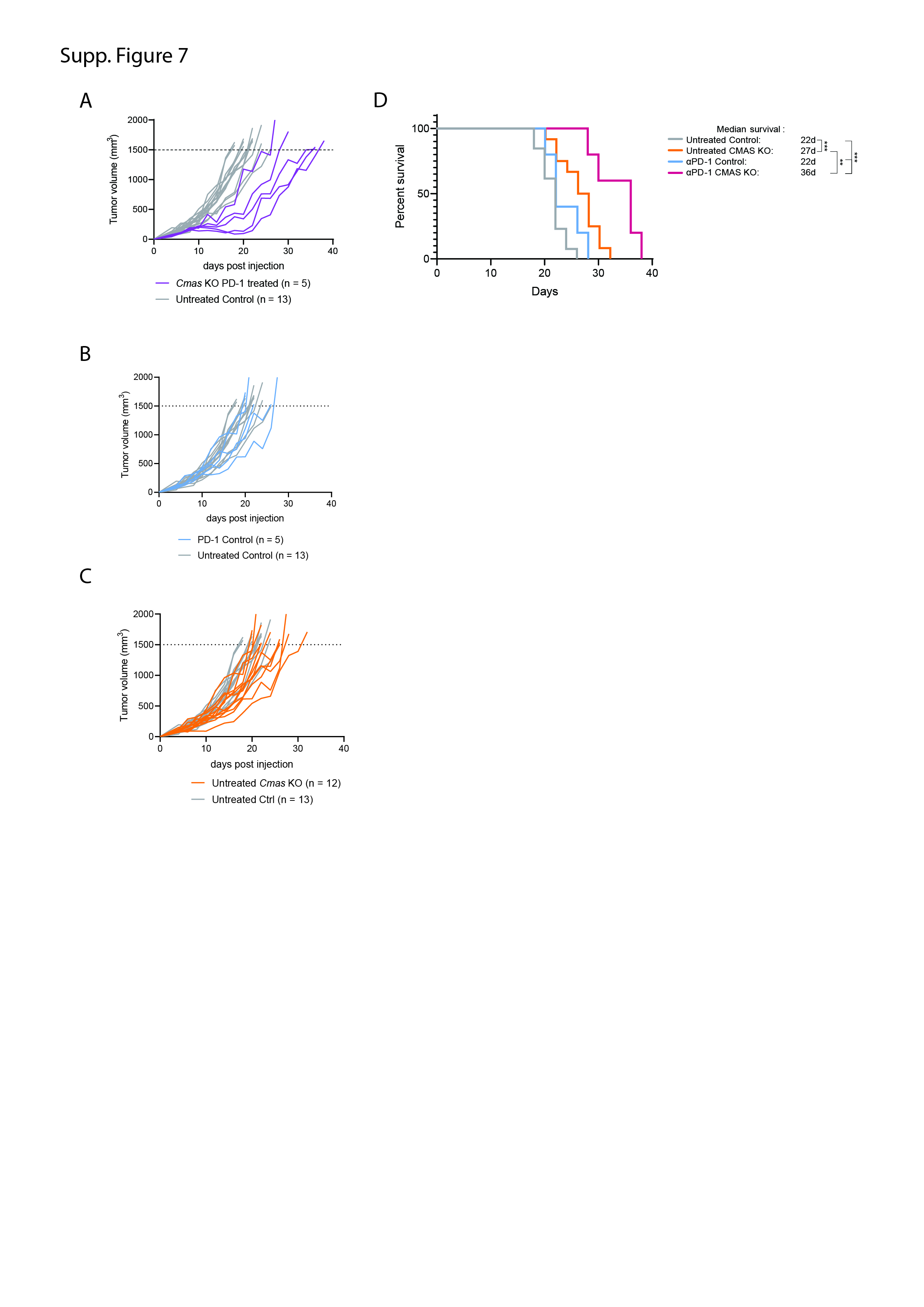

### S Figure 8.jpg

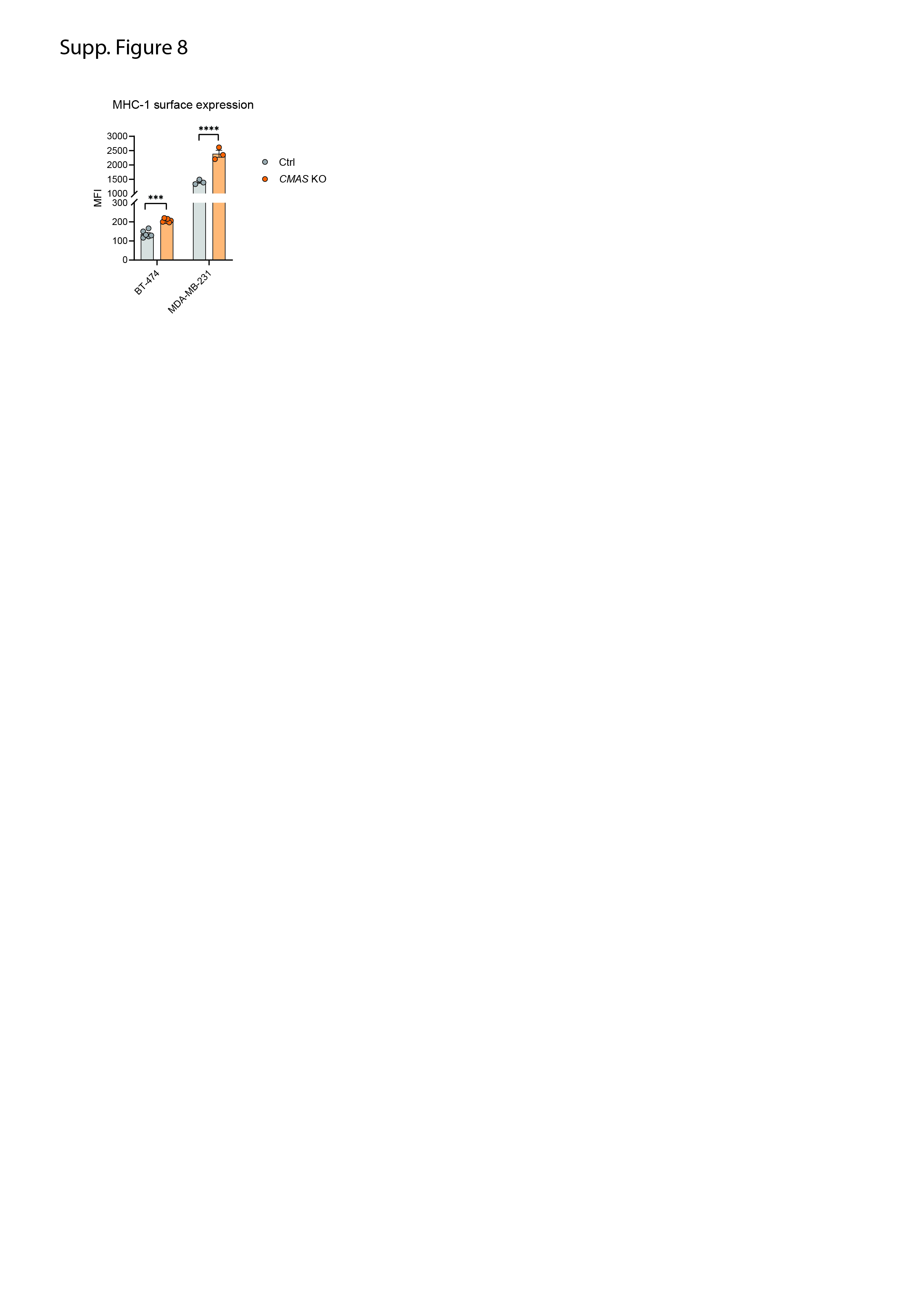

### S Figure 9.jpg

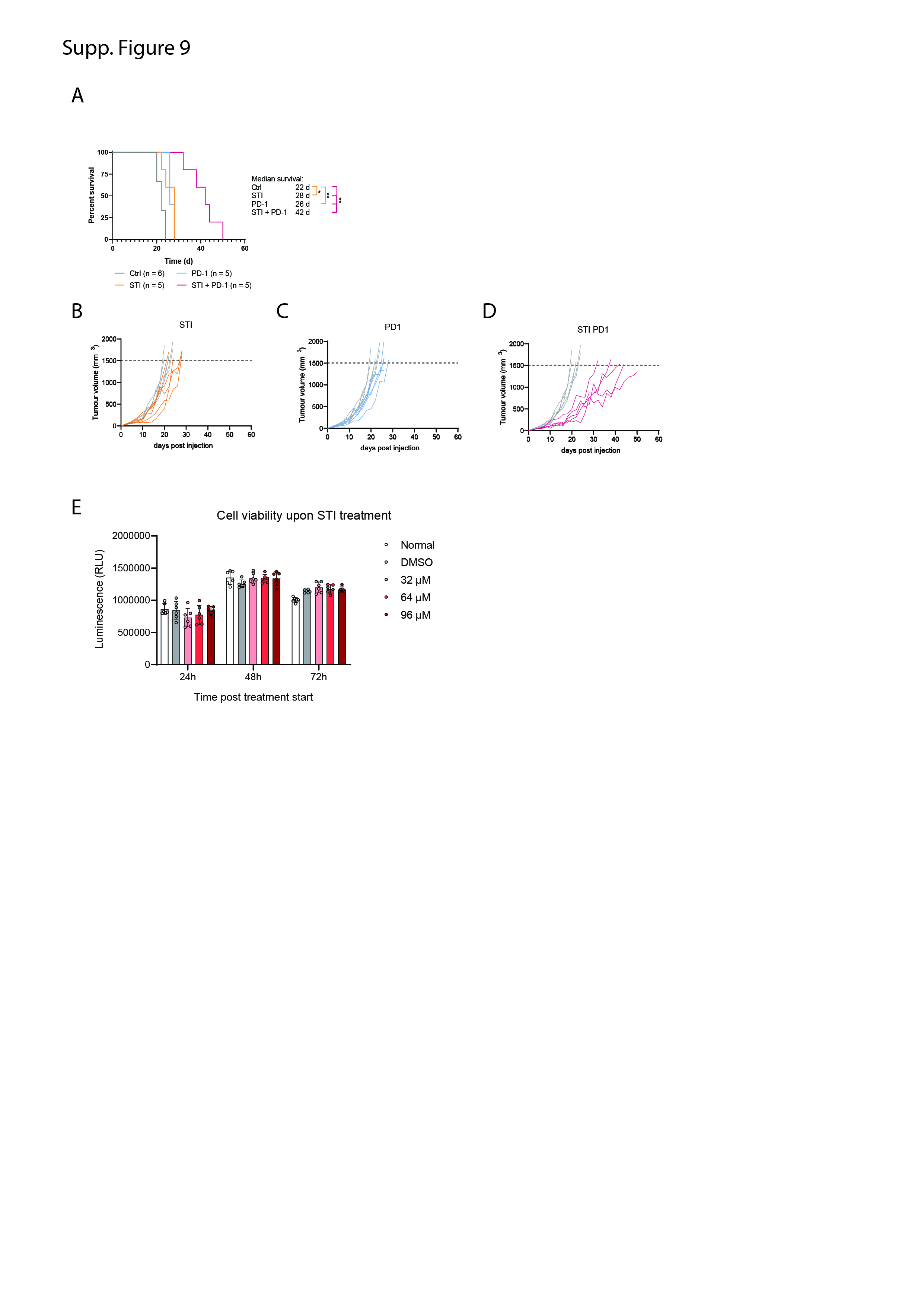

### S Figure 10.jpg

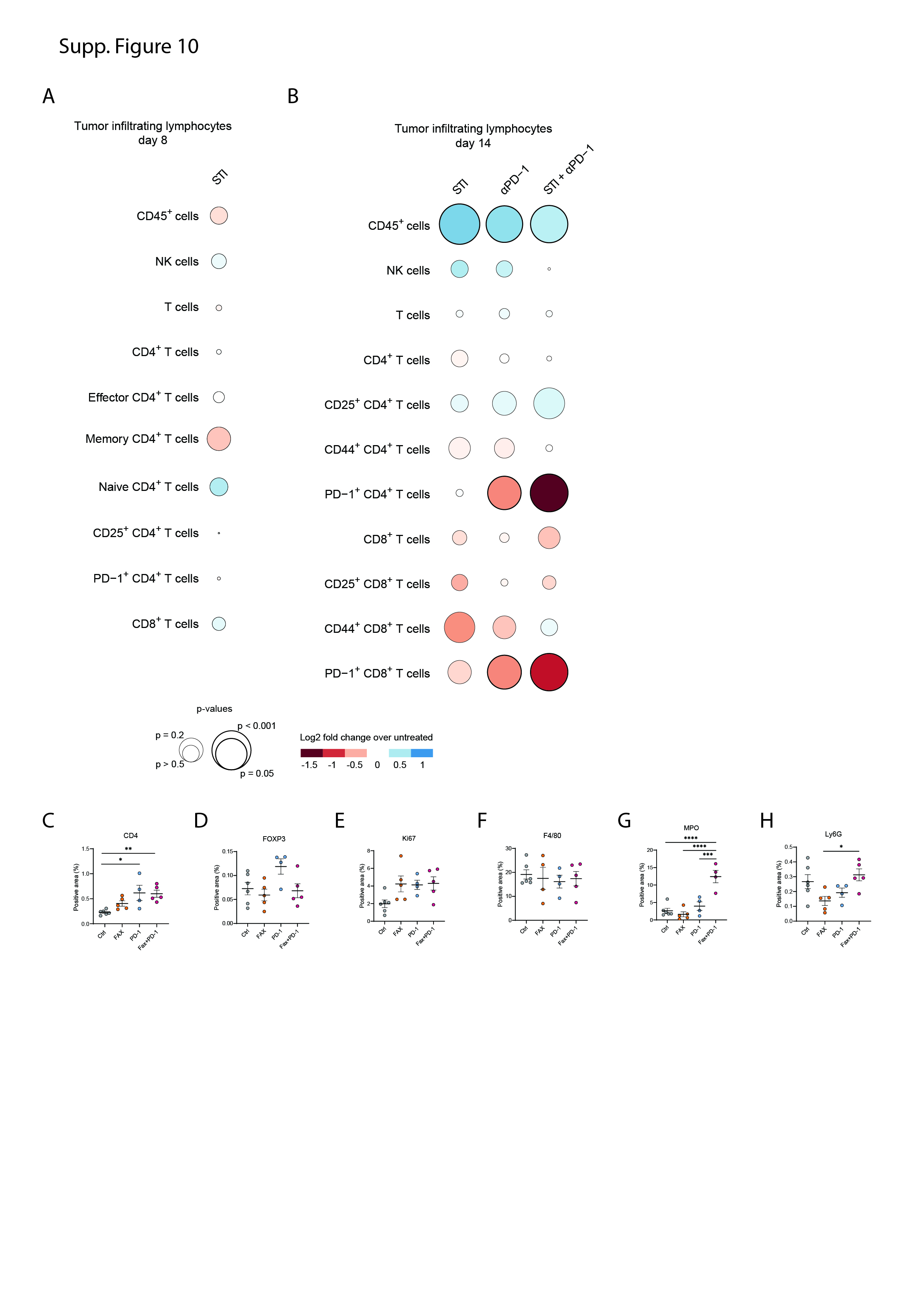

### S Figure 11.jpg

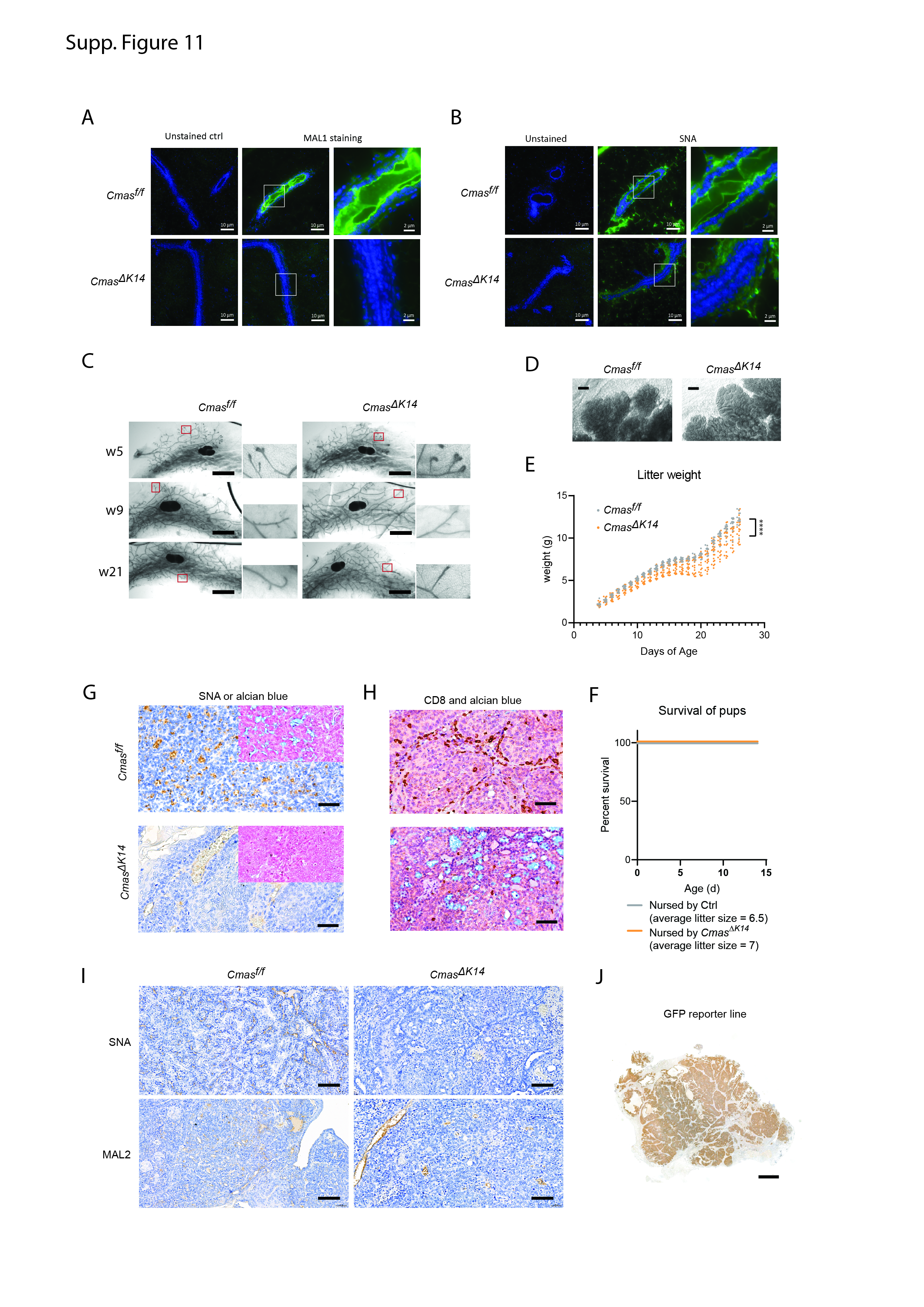

### S Figure 12.jpg

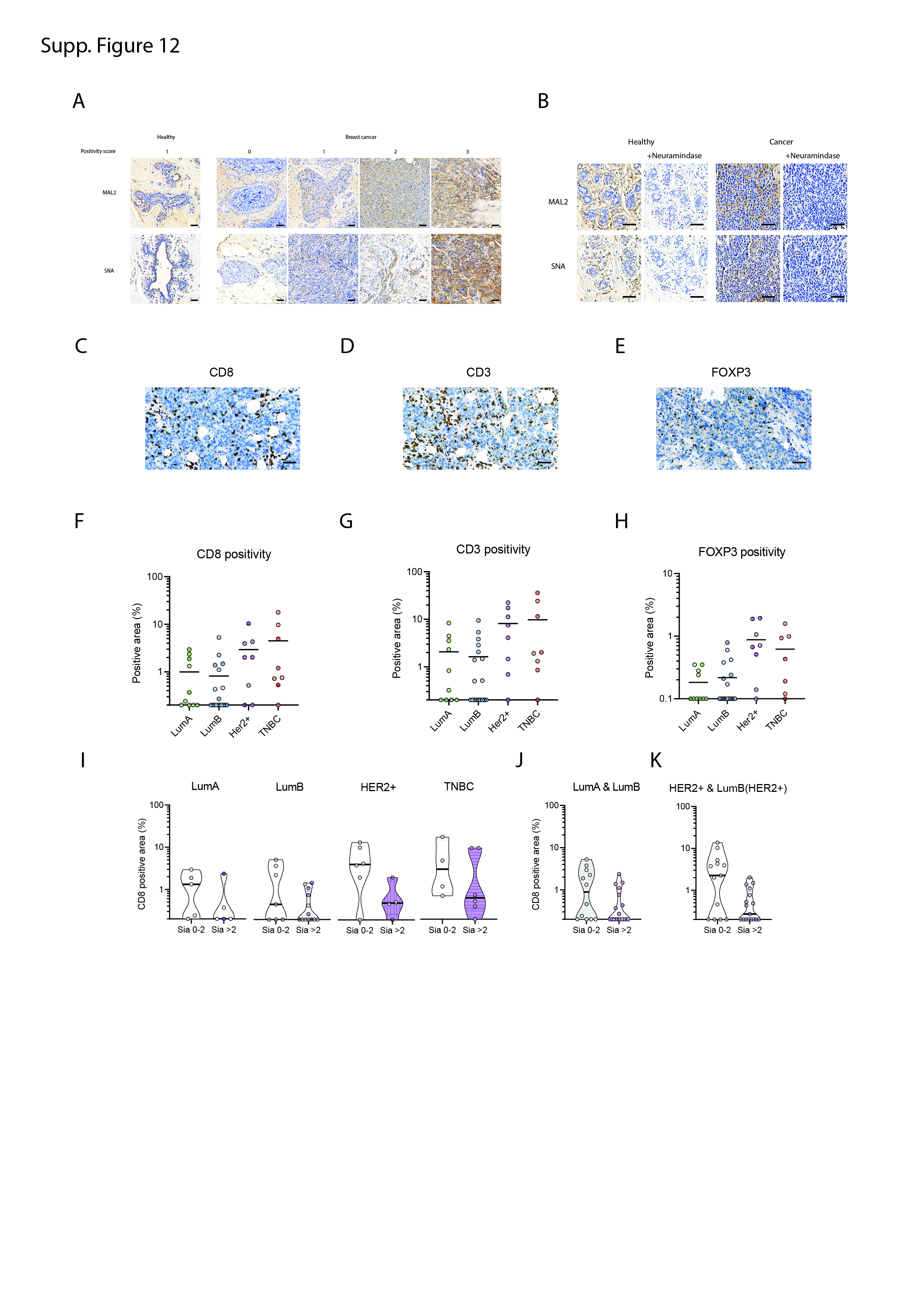
